# Markov modelling of run length and velocity for molecular motors

**DOI:** 10.1101/727636

**Authors:** James L Buchanan, Robert P Gilbert

## Abstract

Molecular motors are nanometer scale proteins involved in various intracellular processes such as cargo transport, muscle contraction and cell division. The steps in the chemo-mechanical cycle that use ATP hydrolysis to generate motion along the cytoskeletal tracks can be characterized by stepping rates depending only on the current state, permitting modeling of the cycle as a Markov process. To learn more about the nature of motor motion, cell researchers have conducted *in vitro* experiments in which molecular motors pull beads along filaments attached to glass slides and their velocity and run length until detachment from the microtubule are recorded. In this article a formula is derived for distance traveled until detachment. Combining this with a standard result for time to absorption for Markov processes gives a formula for velocity. Two kinesins, and two variants of myosin VI, for which there are run length and velocity measurements in the literature are considered. In each case the derived formulas for run length and velocity have a generalized Michaelis-Menten form as functions of ATP concentration. The degrees of the numerator and denominator polynomials increase with the complexity of chemo-mechanical cycle. The degrees of the Michaelis-Menten form determine the maximum number of of reaction rates that can be determined from experimental data on run length and velocity, however consistency conditions may reduce this number and restrict which rates can be determined.

## 1 Introduction

Intracellular cargo is moved along the cytoskeletal tracks by the molecular motors kinesin, dynein, and myosin. Asbury et al. [Asbury et al., 2003], Muthukrishnan et al. [Muthukrishnan et al., 2009], Elting et al. [Elting et al., 2011] and Shastry and Hancock [Shastry and Hancock, 2010] describe *in vitro* experiments in which the procession of molecular motors along microtubule tracks attached to glass plates is measured. The latter three articles give statistics on the length traversed until detachment.

To mathematically model the motion of these molecular motors the rates at which the various steps in the chemo-mechanical ATP hydrolysis cycle occur are needed. The supplement to [Muthukrishnan et al., 2009] indicates that the experimental data presented is not sufficient to determine all of these rates, and thus some must be estimated from experiments described in the literature. However it is then noted that there is a lack of consensus values for these rates. The main objective of this article is to explore what can be inferred from measurements on run length and velocity alone, and thus what must be inferred from other types of experiments.

In [Muthukrishnan et al., 2009] the Gillespie algorithm [Gillespie, 1977] was used as a basis for a Monte-Carlo procedure to simulate the runs of kinesin-1 and -2. In [Elting et al., 2011] fundamental probality considerations were used to predict expected run lengths and run times and a bootstraping technique to determine parameter values. In this article we show that a Markov modeling approach can treat all of the molecular motors.

## 2 Speed and diffusion of molecular motor movement

Elston [Elston, 2000] gives a procedure for finding the coefficients of the diffusion-advection equation

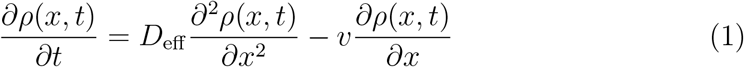

where *ρ* is the probability density function for position along the cytoskeletal track, *D*_eff_ is the diffusivity, and *v* is the advective velocity. The coefficients in (1) are found by considering a Markov process in which *S*(*t*) is the state of the molecular motor and *N* (*t*) its position along the cytoskeletal track at time *t*. Each position *N* is separated by distance Δ*x*. The governing equation for the Markov process is

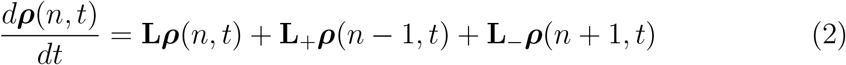

where the *i*th component, *ρ*_*i*_(*n, t*), of the vector ***ρ*** is the probability that *S*(*t*) = *i* and *N* (*t*) = *n*. The formulas given in [Elston, 2000] for velocity and diffusivity in (1) are

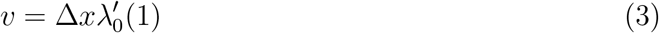

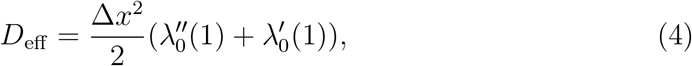

where *λ*_0_(*z*) is the largest eigenvlaue of the matrix

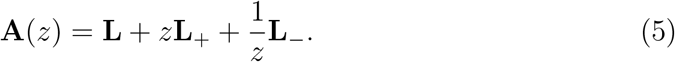

## 3 Expected run length and run time for a simple three-stage model

The formulation in [Elston, 2000] can be adapted to predict run length and its dispersion for the type of experiments described in [Asbury et al., 2003], [Muthukrishnan et al., 2009], [Elting et al., 2011] and [Shastry and Hancock, 2010]. In Appendix A it is shown that the expected run length is given by (A.7)

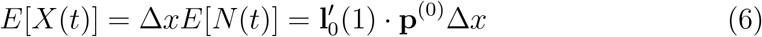

where **l**_0_(1) is a normalized left eigenvector of *A*(1) corresponding to the largest eigenvalue *λ*_0_(1) and **p**^(0)^ = (*p*_0_, *p*_1_, …, *p*_*M*_) is the vector of the probabilities for the starting state. The expected run time given by (A.9) with *j* = 1,

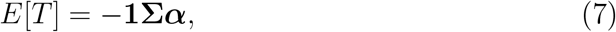

where **1** is a row vector of 1’s, 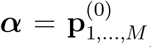 and the subgenerator matrix for this problem is **Σ** = *A*(1)_1,…,*M*,1,…,*M*_ is a standard result on time to absorption for continuous-time Markov processes.

As an illustration Figure 1 shows a model consisting of four states: (0) both heads detached; (1) both heads attached; (2) front head unattached; (3) rear head unattached. It is assumed that the detachment rates of the heads in state (1) are different for front and back heads, perhaps due to tension effects of the linker between the two heads, but that the rate of detachment, *k*_Detach_, and reattachment, *k*_Attach_, are the same for states (2) and (3). The matrix (5) is

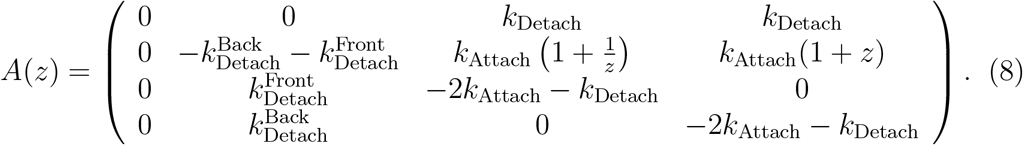

**Figure 1:**
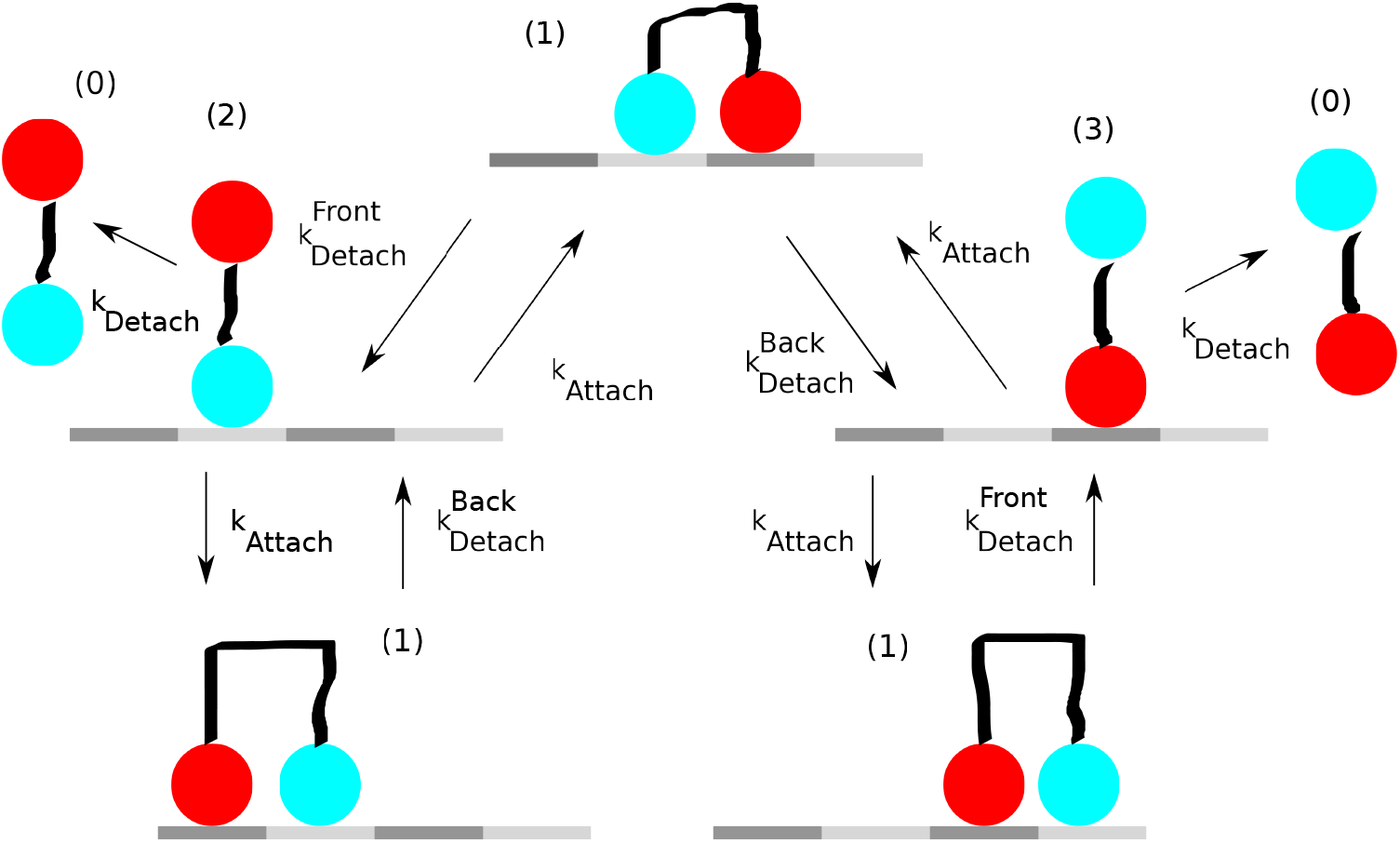
Simple model of a molecular motor with forward and backward stepping.

The diagonal entries are chosen to make each column of *A*(1) sum to zero.

From (6), the expected number of steps, 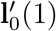, starting from states (0)-(3) is

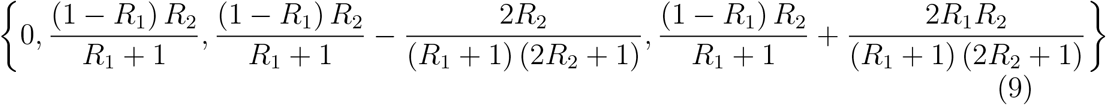

where 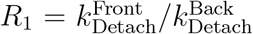 and 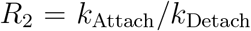. If *R*_1_ = 0, that is, the front head cannot detach in state (1) and thus the motion is monotonically forward, then the expected number of steps is *R*_2_. This can be derived directly from elementary considerations involving the duty ratio ([Elting et al., 2011]).

From (7), the expected time to separation starting from state (1) is

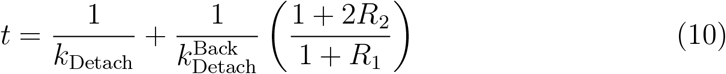

and thus the steps per second are

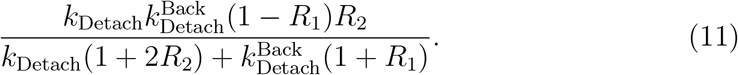

Figure 2 shows the expected number of steps as a function of *R*_2_ when *R*_1_ = 0.5. Figure 3 shows the expected number of steps per second when *R*_1_ = 0, 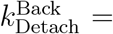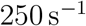, *k*_Detach_ = 1.7 s^−1^ (cf Table 1).

**Table 1:** Parameters for the kinesin-1 model of Figure 4. From [Muthukrishnan et al., 2009].

| Rate constant | Forward | Abbrev | No. | Backward | Abbrev | No. |
| --- | --- | --- | --- | --- | --- | --- |
| $k_{\text{detach}}$ | $250 \text{ s}^{-1}$ | $k_{12}$ | 1 | $0.1 \text{ s}^{-1}$ | $k_{21}$ | 2 |
| $k_{\text{onATP}}$ | $2 \mu\text{M}^{-1} \text{ s}^{-1}$ | $\tilde{k}_{23}$ | 3 | $200 \text{ s}^{-1}$ | $k_{32}$ | 4 |
| $k_{\text{hydrolysis}}$ | $280 \text{ s}^{-1}$ | $k_{34}$ | 5 | $3.5 \text{ s}^{-1}$ | $k_{43}$ | 6 |
| $k_{\text{attach}}$ | $370 \text{ s}^{-1}$ | $k_{41}$ | 7 | $0.1 \text{ s}^{-1}$ | $k_{14}$ | 8 |
| $k_{\text{unbind2}}$ | $0.003 \text{ s}^{-1}$ | $k_{20}$ | 9 | | | |
| $k_{\text{unbind}}$ | $1.7 \text{ s}^{-1}$ | $k_{40}$ | 10 | | | |

**Figure 2:**
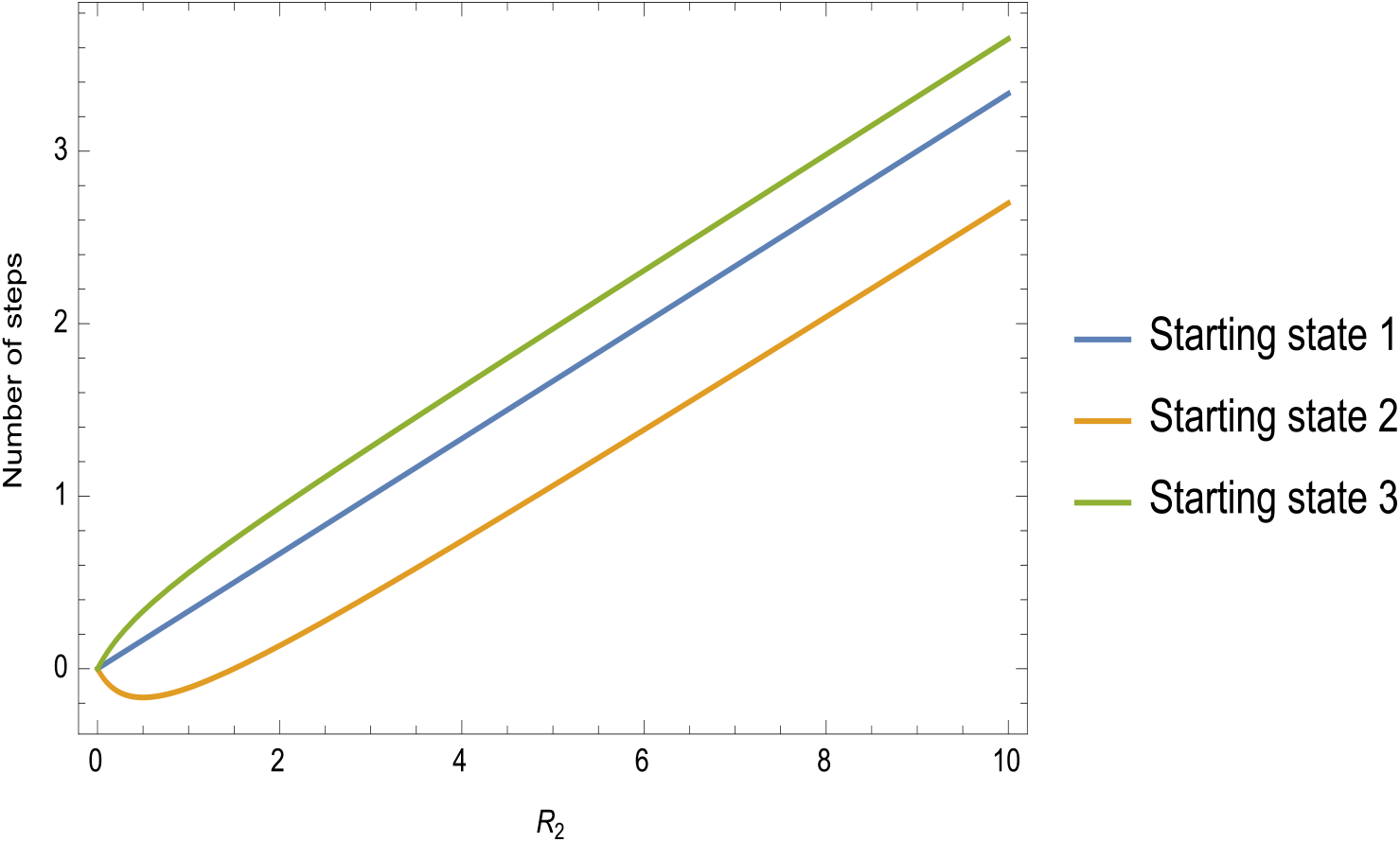
Expected number of steps from different starting states in Figure 1 when *R*_1_ = 0.5.

**Figure 3:**
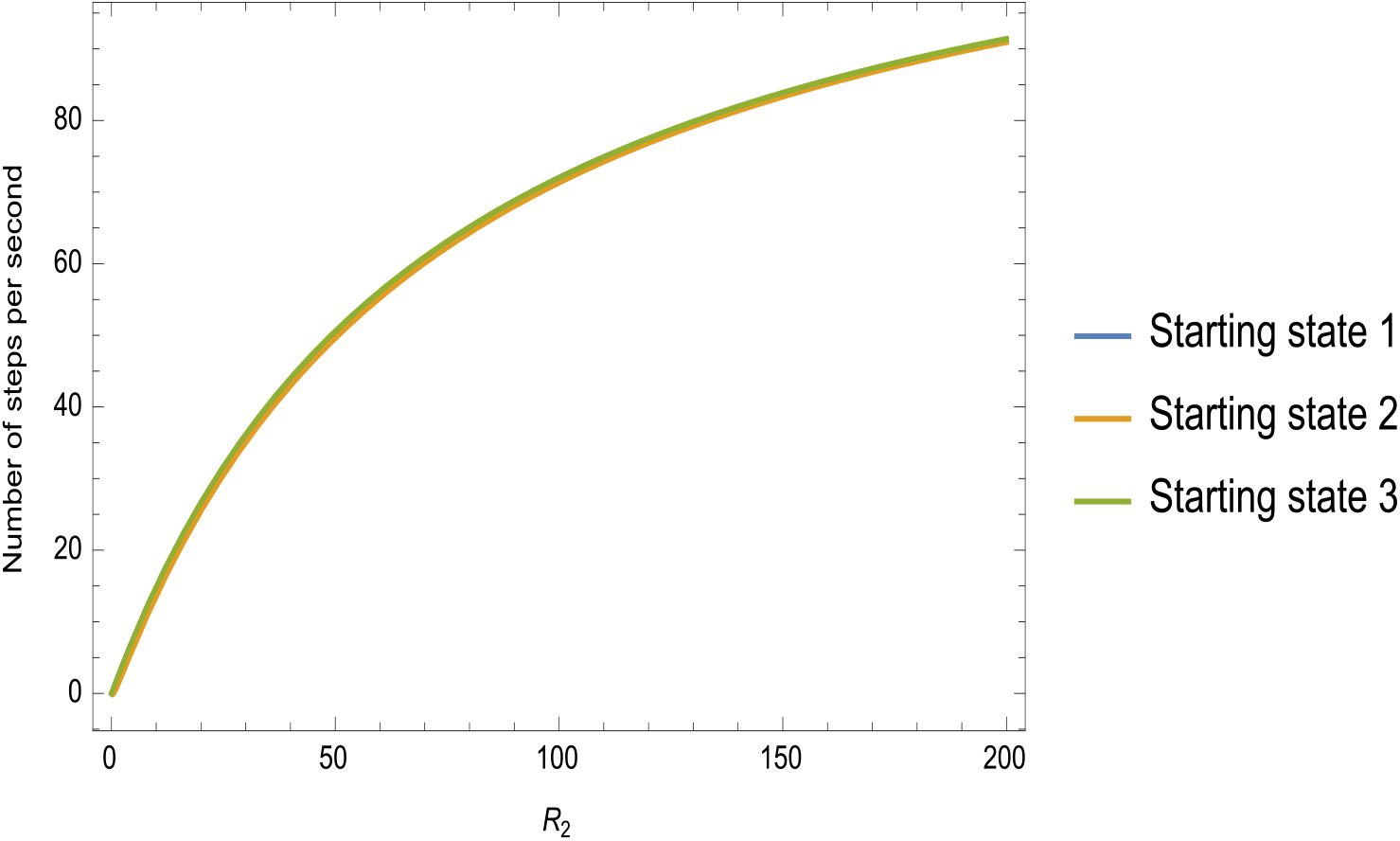
Expected number of steps per second from different starting states in Figure 1 when *R*_1_ = 0, 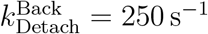 *, k*_Detach_ = 1.7 s^−1^.

**Figure 4:**
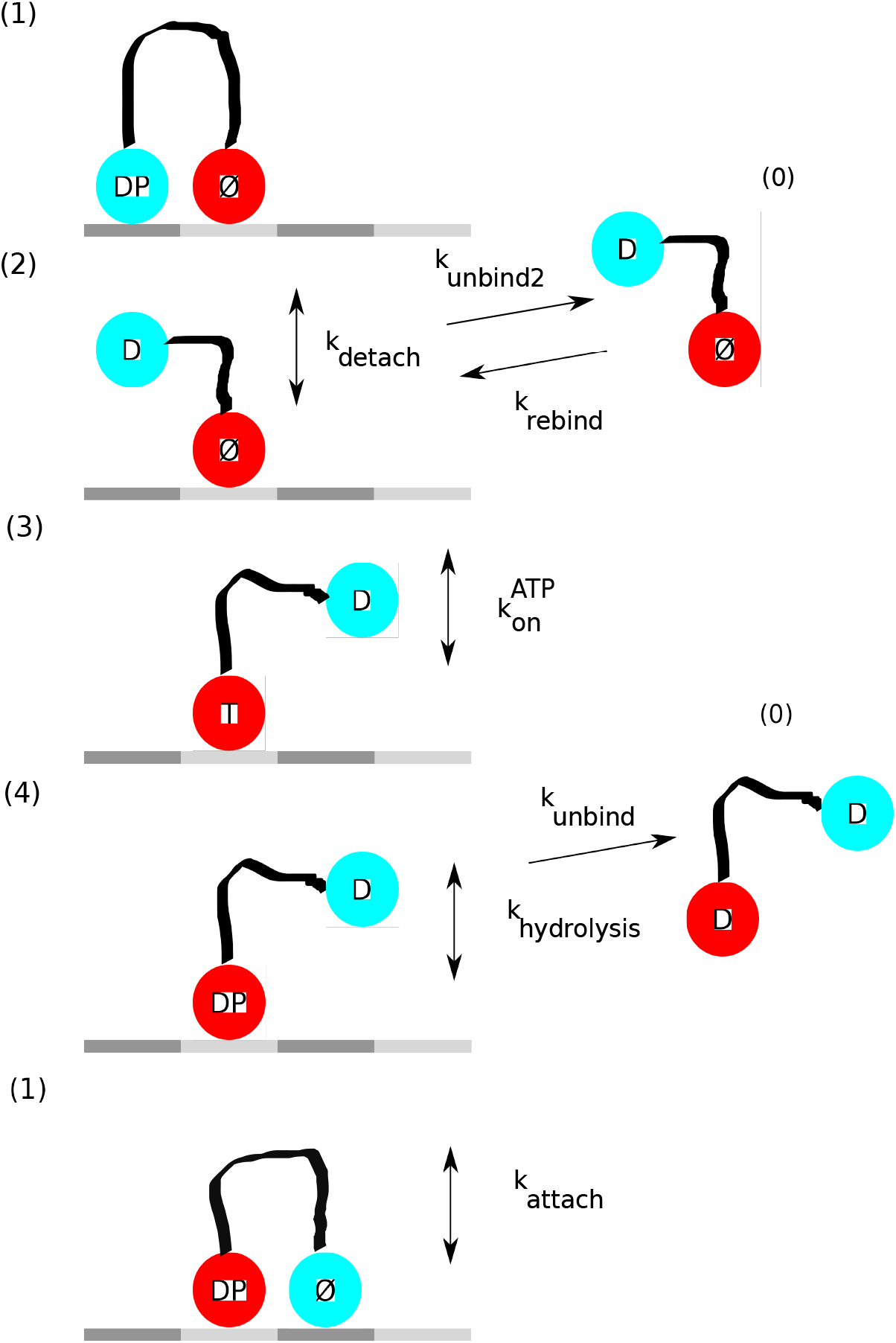
After Figure 1 from [Muthukrishnan et al., 2009]. D: ADP; T: ATP; P: Phosphate; *∅*: No nucleotide.

## 4 Expected run length and run time for kinesin-1: Michaelis-Menten formulation

The article [Muthukrishnan et al., 2009] presents measurements of run length and velocity of kinesin-1 and 2. For the model for kinesin-1 shown in Figure 4, the matrix (5) becomes

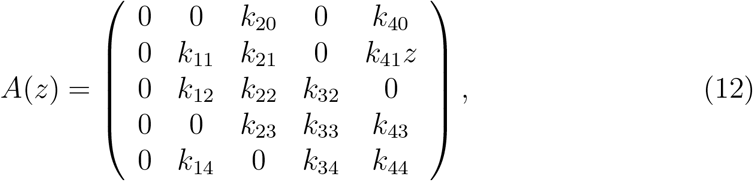

if the possibility of backward steping is neglected. As indicated in Section 3, allowing for backward steppping would require roughly doubling the number of states. Here *k*_*ij*_ denotes the stepping rate from state *i* to state *j*. Thus from Figure 4,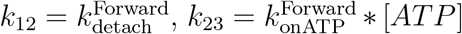, where [*ATP*] is the concentration of ATP in *μM*, *k*_40_ = *k*_unbind_, and so forth. The possibility of rebinding, *k*_02_ = *k*_rebind_, is neglected.

When run length is computed from (6) and velocity is computed as the quotient of run length (6) and run time (7) (as opposed to using (3), the results have Michaelis-Menten (1:1) form

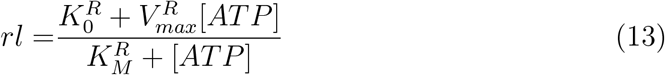

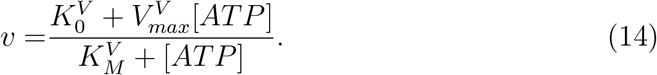

where

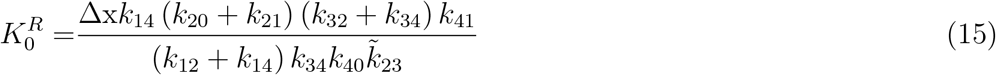

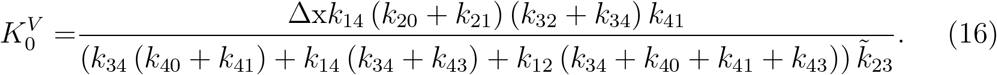

The coefficients 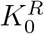 and 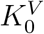 are small, 0.06 and 0.03 respectively, relative to the other numerator term for the values of the parameters in Table 1, unless [ATP] is much less than 1 *μ*M. They vanish if *k*_14_ = 0 in which case run length and velocity have the standard Michaelis-Menten form with

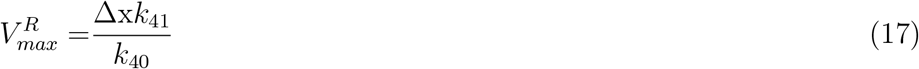

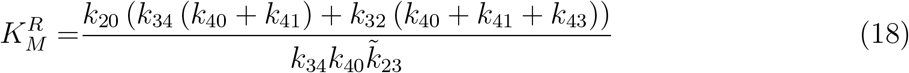

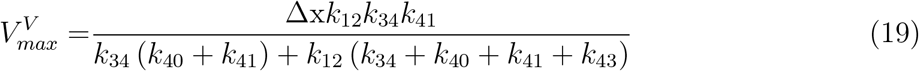

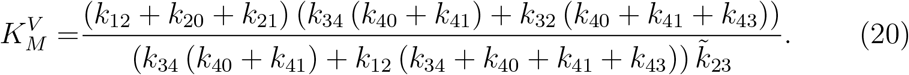

In the above formulas 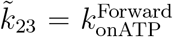. The only occurence of *k*_21_ is now in a sum with *k*_12_, which is expected to be much larger. Thus *k*_21_ = 0 will also be assumed.

It is noted in [Muthukrishnan et al., 2009], Figure 3, that the experimental data for velocity for kinsein-1 fits Michaelis-Menten form well with 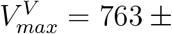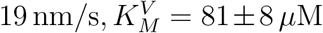. The parameter values in Table 1 give 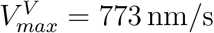, 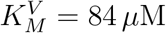 when used in (19) and (20).

Figure 5, left panes, compares the predictions of (13) and (14) to the experimental measurements shown in Figure 3 of [Muthukrishnan et al., 2009]. In the absence of the possibility of detachment from state 2 of Figure 4, *k*_20_ = 0, the run length is predicted to be 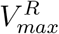, independent of ATP concentration. This near independence was observed in experiments by [Yajima et al., 2002]. However [Muthukrishnan et al., 2009] found that the experimental measurements for run length shown in Figure 5, upper left pane, only narrowly missed, *p* = 0.07, the conventional threshold, *p* = 0.05, for being significantly different from constant.

**Figure 5:**
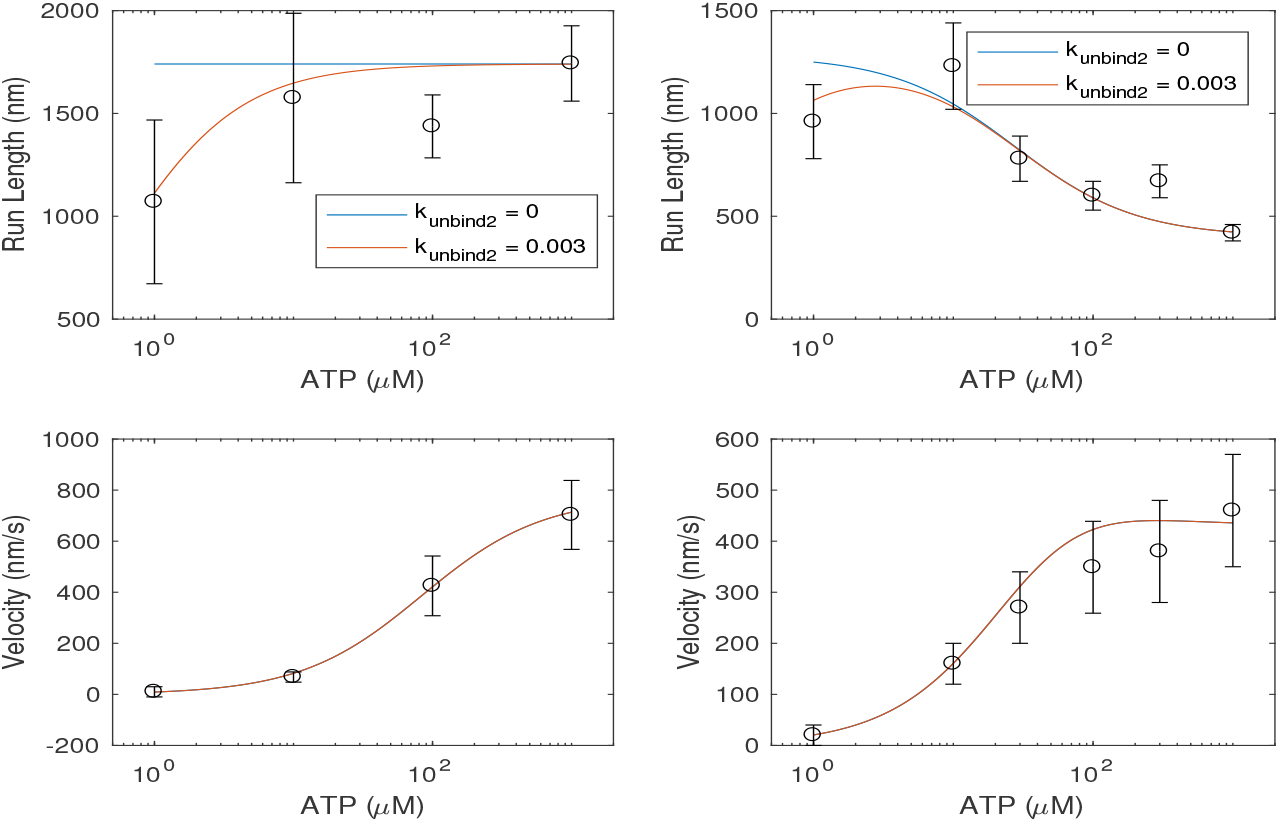
Markov model predictions for kinesin-1 (left panes) and kinesin-2 (right panes) when the parameters used are those of Tables 1 and 3 respectively. The data points and the standard error (SE) estimates were read from Figure 3 of [Muthukrishnan et al., 2009]. The step length for both kinesins was taken to be 8 nm.

Since the formulas (6) and (A.8) are asymptotic in time, confirmation of their accuracy was sought by comparing them to the averaged results of 5000 simulations of the Markov process with state changes being determined pseudorandomly using the formula

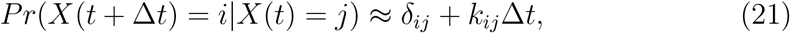

and simply tracking the positions of the two heads. Thus all that is assumed is the fundamental Markov hypothesis that the probability of moving to the next state depends only upon the current state.

For the parameters in Table 1 the expected run length and velocity were well-predicted by (6) and (7) in two cases: 1) the parameters are as given in Table 1, 2) *k*_unbind2_ = 0 with the remaining parameters as in Table 1. It also should be noted that no instances of full backward steps were encountered in the 5000 trials unless large, order of 10*s*^−1^, backward rates were used. Thus the assumption *k*_14_ = 0, which precludes backward steps, should introduce little distortion of the model’s predictions.

**Table 2:**
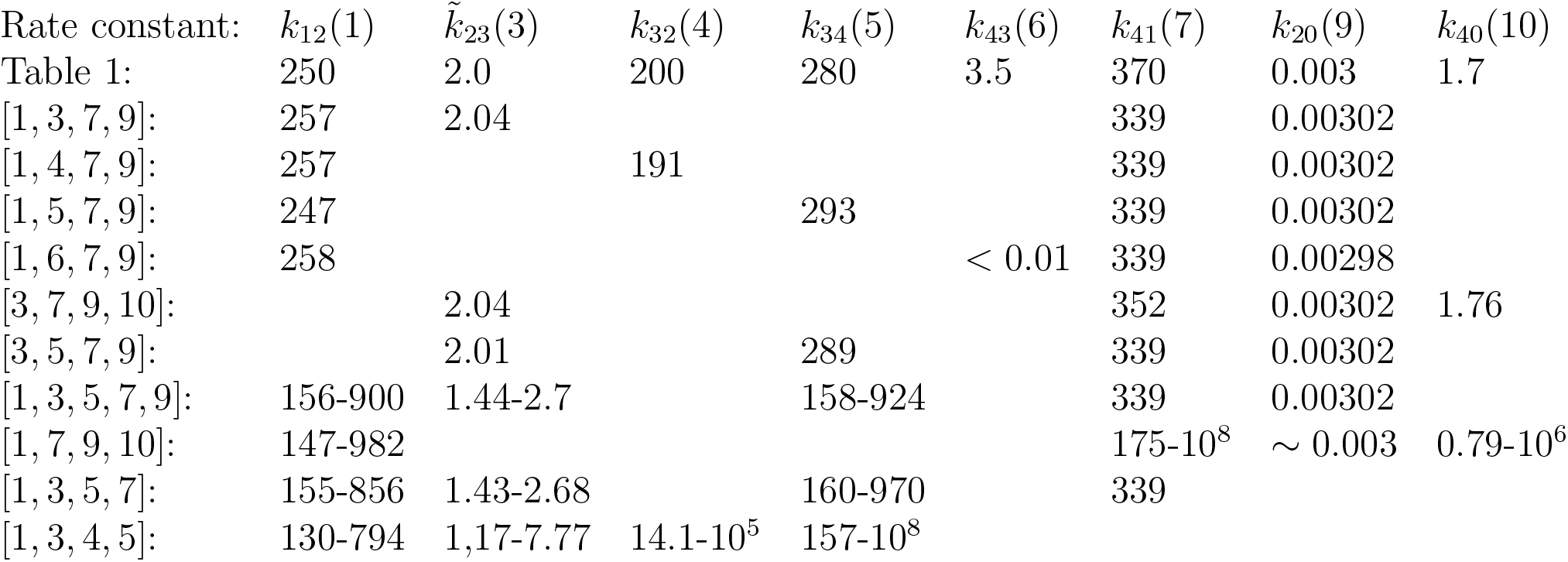
Numerical determination of some the kinesin-1 parameters by minimiz-ing the discrepancy with experimental measurements. Ten trials were run with randomly chosen initial guesses for the parameters based on their order of magni-tude. If more than four parameters were sought, the ten trials resulted in a wide range of values with about the same objective function value, as illustrated in the [1, 3, 5, 7, 9] entry. This was also sometimes the case when four parameters were sought.

**Table 3:** Parameters for the kinesin-2 model of Figure 12. From [Muthukrishnan et al., 2009].

| Rate constant | Forward | Backward |
| --- | --- | --- |
| $k_{\text{detach}}$ | $250 \text{ s}^{-1}$ | $0.1 \text{ s}^{-1}$ |
| $k_{\text{onATP}}$ | $3 \mu\text{M}^{-1} \text{ s}^{-1}$ | $15 \text{ s}^{-1}$ |
| $k_{\text{hydrolysis}}$ | $120 \text{ s}^{-1}$ | $3.5 \text{ s}^{-1}$ |
| $k_{\text{attach}}$ | $370 \text{ s}^{-1}$ | $0.1 \text{ s}^{-1}$ |
| $k_{\text{fhdettach}}$ | $11 \text{ s}^{-1}$ | $0 \text{ s}^{-1}$ |
| $k_{\text{unbind}}$ | $2.3 \text{ s}^{-1}$ | |
| $k_{\text{unbind2}}$ | $0.003 \text{ s}^{-1}$ | |

Another series of numerical experiments was to attempt to determine numerically some of the parameters of the kinesin-1 model of Figure 4. MATLAB’s constrained minimizer *fmincon* was used to find the values of various subsets of the parameters of the model, with the remaining parameters as given in Table 1. The objective function minimized the *l*_2_ norm of the vector of relative errors between the calculated run length and velocity and the experimental measurements shown in Figure 5. Ten trials were run with initial guesses chosen by a random number generator based only on the expected order of magnitude of the parameter, which was assumed known. The values of the parameters were constrained to be positive. Table 2 shows that the minimization process obtained definite and reasonable values for subsets of at most four parameters. This is consistent with the derived two parameter Michaelis-Menten forms for run length and velocity. The minimizations were not always successful for subsets of four. This will be explored further.

Since the four experimental data points for run length shown in Figure 5 are not monotonically increasing with ATP concentration, and therefore not compatible with the derived Michaelis-Menten form, Mathematica’s *FindFit* was used to fit curves to subsets of the data points. The results are shown in Figure 6. The curve fit for all four data points came close to lying within the standard error estimates, as did the curve fit that omitted the fourth, rightmost, data point. Figure 7 shows that omitting the rightmost data point does not produce good agreement with the velocity data.

**Figure 6:**
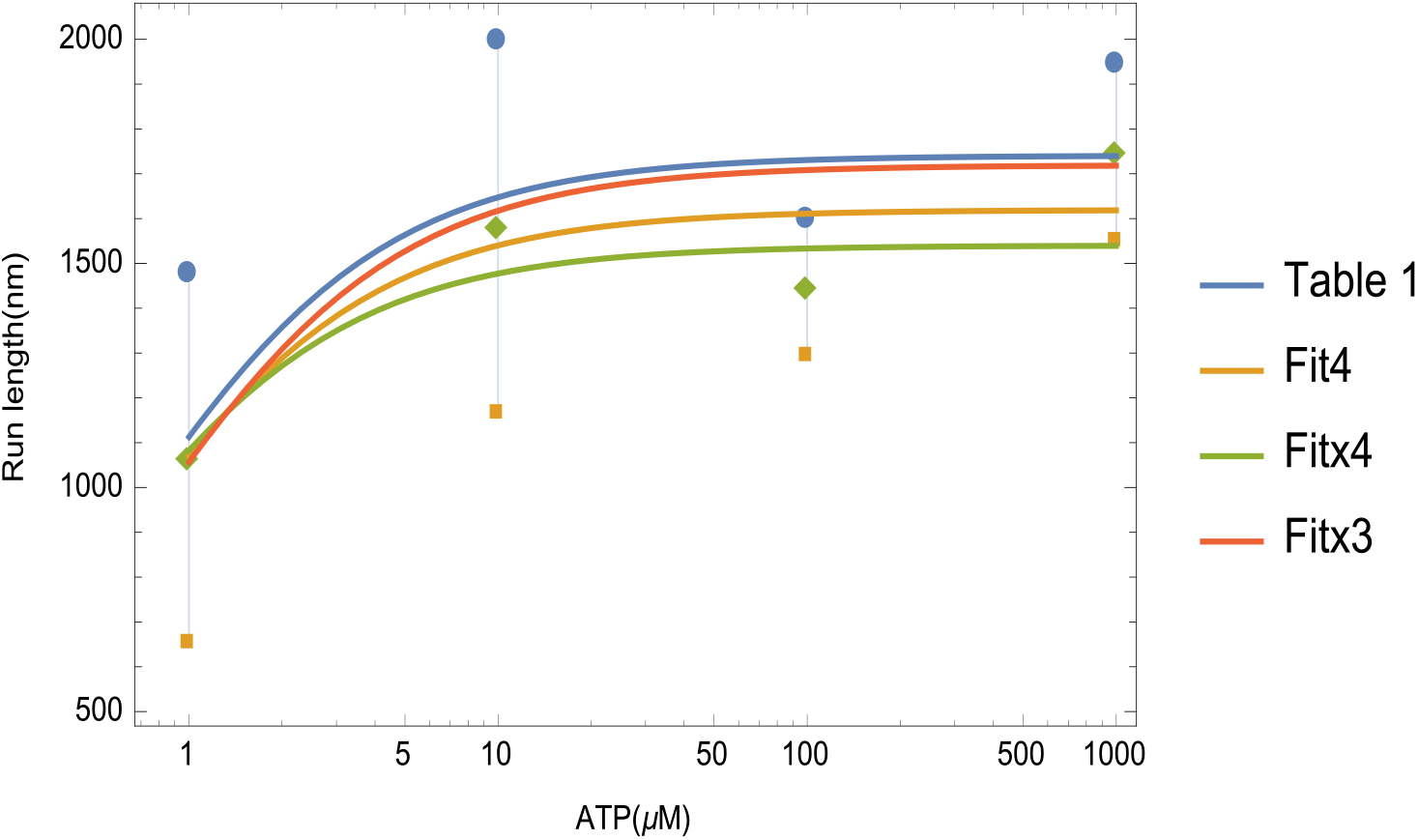
Four determinations of the Michaelis-Menten parameters from the ex-perimental data for run length for kinesin-1. Table 1: Michaelis-Menten curve for the parameters given in Table 1; Fit4: Curve fit of all four data points; Fitx3: Curve fit of data points 1,2,4; Fitx4: Curve fit of data points 1,2,3.

**Figure 7:**
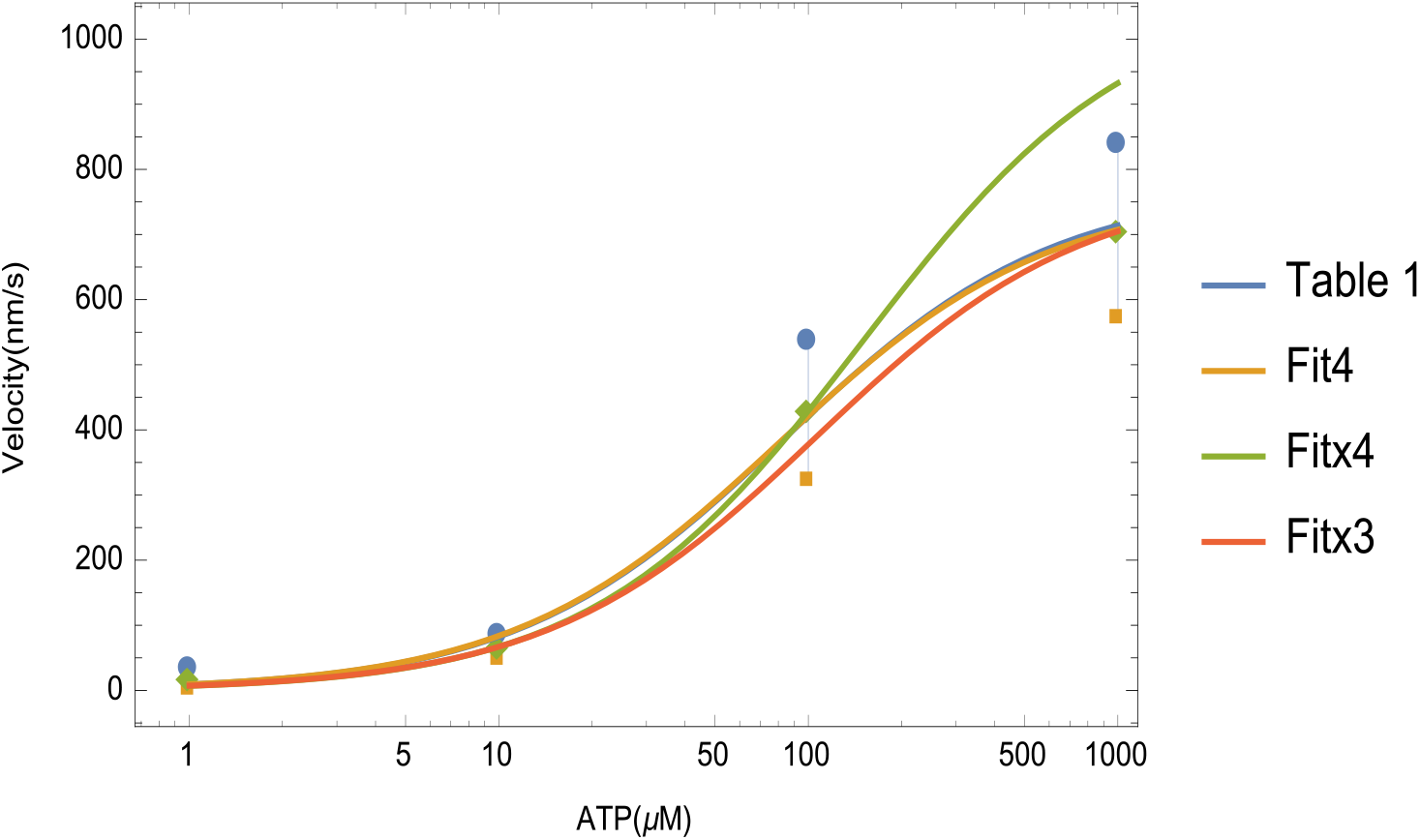
Four determinations of the Michaelis-Menten parameters from the ex-perimental data for velocity for kinesin-1. Table 1: Michaelis-Menten curve for the parameters given in Table 1; Fit4: Curve fit of all four data points; Fitx3: Curve fit of data points 1,2,4; Fitx4: Curve fit of data points 1,2,3.

Figures 8 and 9 give two determinations of subsets of four parameters, {*k*_12_, *k*_32_*, k*_41_*, k*_20_ and 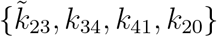 which Table 2 indicates were found successfully using constrained minimization. The parameters *k*_12_*, k*_32_*, k*_41_*, k*_20_ of Figure 8 are those which were determined by [Muthukrishnan et al., 2009] empirically from the experimental data (Table S2), rather than from the literature. To get a notion of the sensitivity to error, the parameter 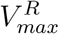 was varied over the range 1500 to 1900 indicated in Figure 6 as the range of run lengths at ATP concentration 1000 *μ*M, with 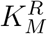 being chosen to make the Michaelis-Menten curve (13) pass through the upper, lower, or midpoint of the error range at ATP concentration 1 *μ*M. Parameters other than the four solved for were taken from Table 1. The two figures indicate that only *k*_20_ = *k*_unbind2_ is sensitive to the measurement of run length at an ATP concentration of 1 *μ*M. The other three parameters showed only moderate sensitivity to the variation of the run length measurement at [ATP]=1000 *μ*M over its standard error range.

**Figure 8:**
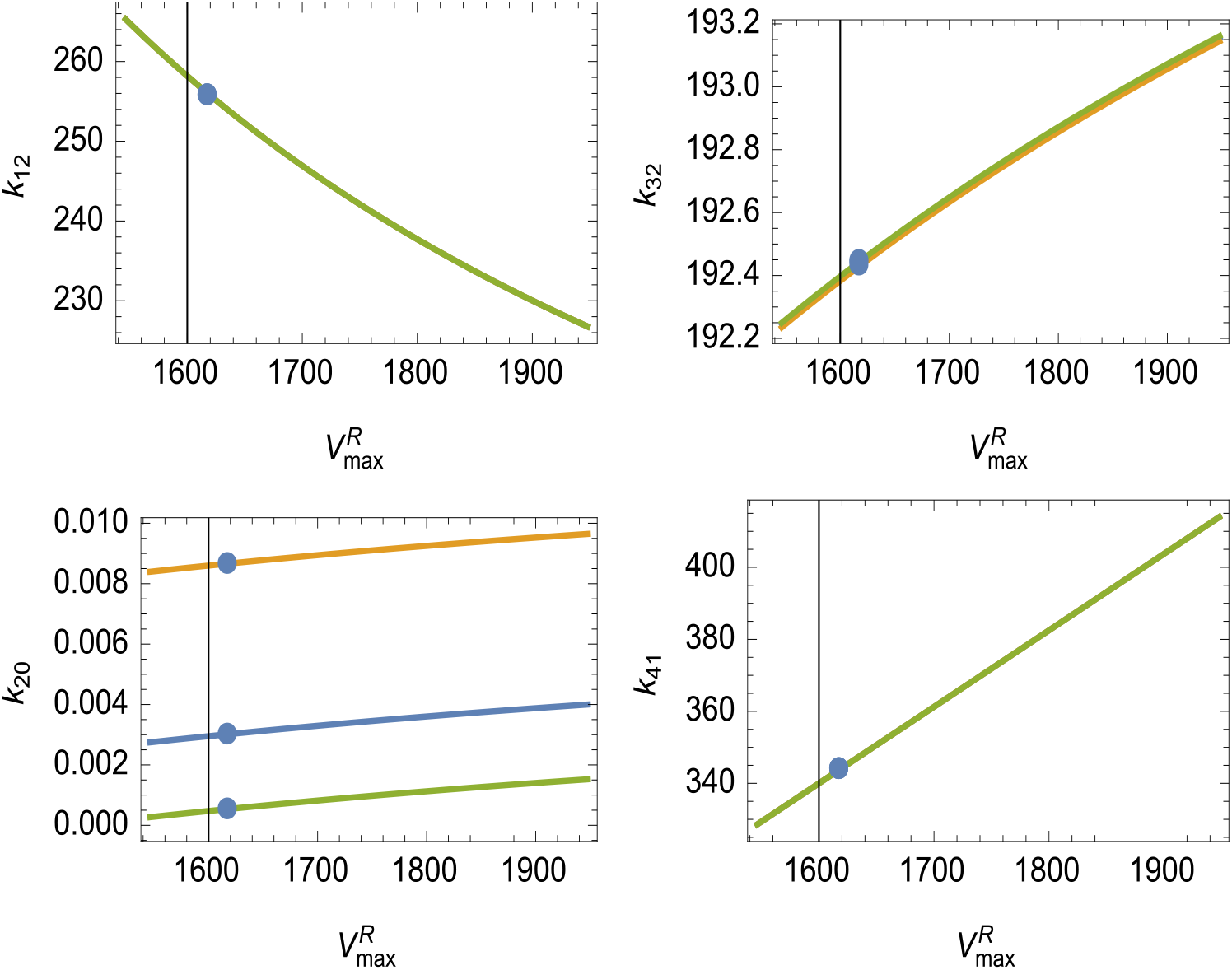
Sensitivity to the parameter 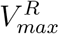 when 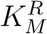 is chosen to make the Michaelis-Menten curve pass through the high (Green), low (Orange) or mid (Blue) point of the experimental error range at 1*μM* of ATP. All other parameter values were taken from Table 1. Blue dots are the parameter values corresponding to “Fit4” in Figure 6.

**Figure 9:**
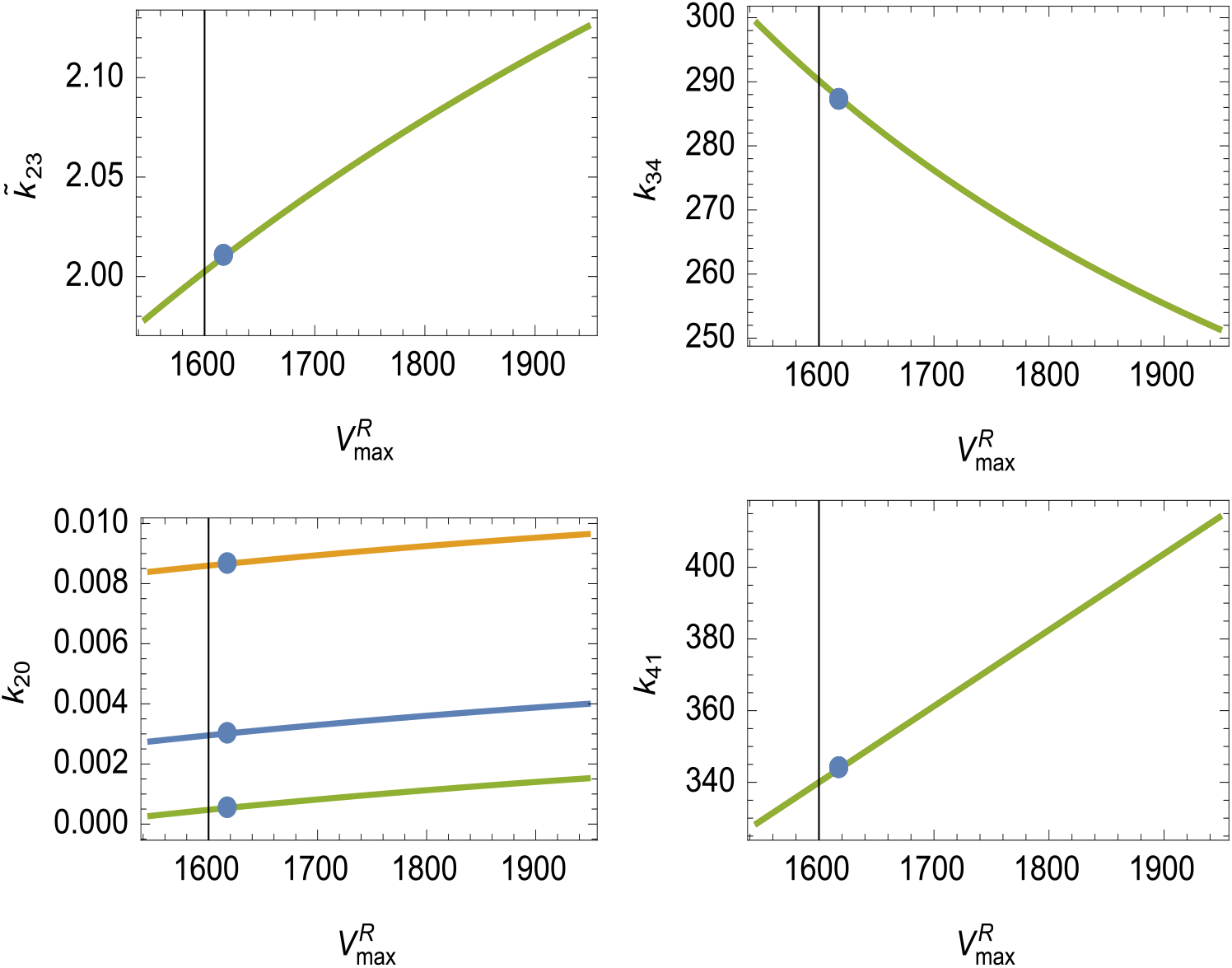
Sensitivity to the parameter 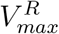 when 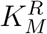 is chosen to make the Michaelis-Menten curve pass through the high (Green), low (Orange) or mid (Blue) point of the experimental error range at 1*μM* of ATP. All other parameter values were taken from Table 1. Blue dots are the parameter values corresponding to “Fit4” in Figure 6.

Since there are 70 ways to choose subsets of four from the eight available parameters it is tedious to try to catalog all of the possible outcomes. To get to fewer parameter sets, some added assumptions are useful. An estimate of the Michaelis-Menten parameters for the dependence of *k*_40_ on ATP concentration can found in [Hancock and Howard, 1999], Figure 2. If *k*_40_ is treated as known, then (17) determines *k*_41_, and the remaining three equations (18)-(20) can be used to find three other parameters. The sets 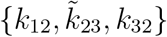 and {*k*_12_, *k*_34_, *k*_43_} cannot be solved for using these the three equations. Solving for *k*_12_ and any of the three pairs 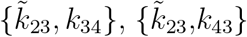, or {*k*_32_, *k*_34_} lead to

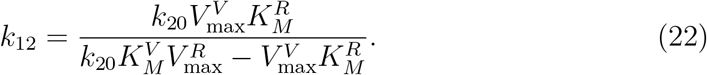

Figure 10 shows that this equation has a vertical asymptote which makes determination of *k*_12_ from a determination of *k*_20_ from some other experiment effectively impossible. Thus if *k*_12_ is to be determined from run length and velocity measure-ments, *k*_20_ must also be determined from these measurements, as it was in the first four cases in Table 2.

**Figure 10:**
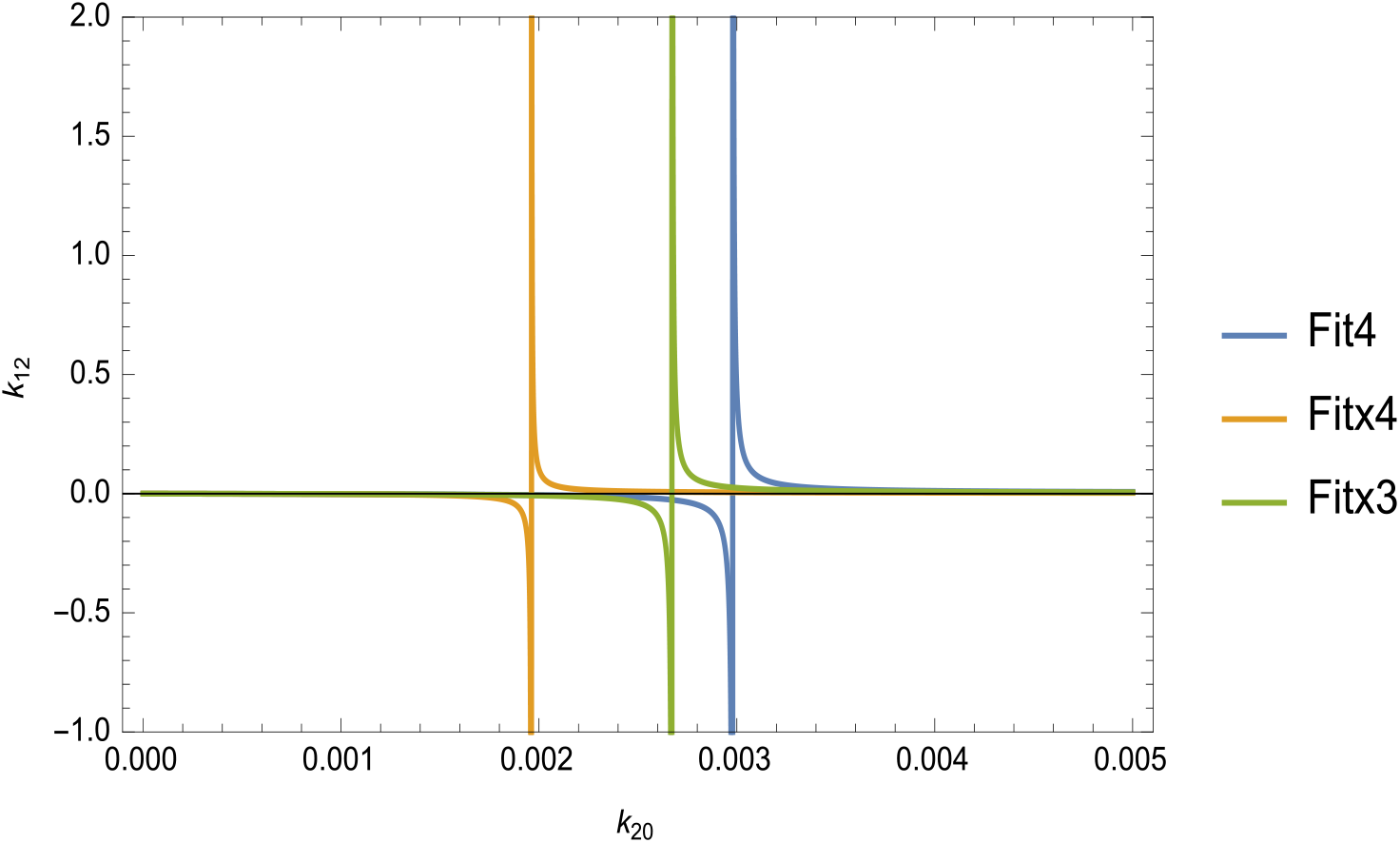
Attempts to solve for *k*_41_, *k*_12_ and any of three pairs of parameters given in the text lead to equation (22). Thus accurate determination of *k*_12_ is precluded, absent extremely accurate determination of *k*_20_.

Other reasons for some of the failures listed in Table 2 are posssible. Figure 11 shows that in solving for {*k*_12_, *k*_20_, *k*_41_, *k*_40_}, the parameters *k*_41_ and *k*_40_ are both negative throughout the range of 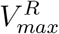 under consideration. The detachment parameter *k*_12_ is positive, but less than the ∼ 100 s^−1^ minimum implied by the overall speed of the motor protein.

**Figure 11:**
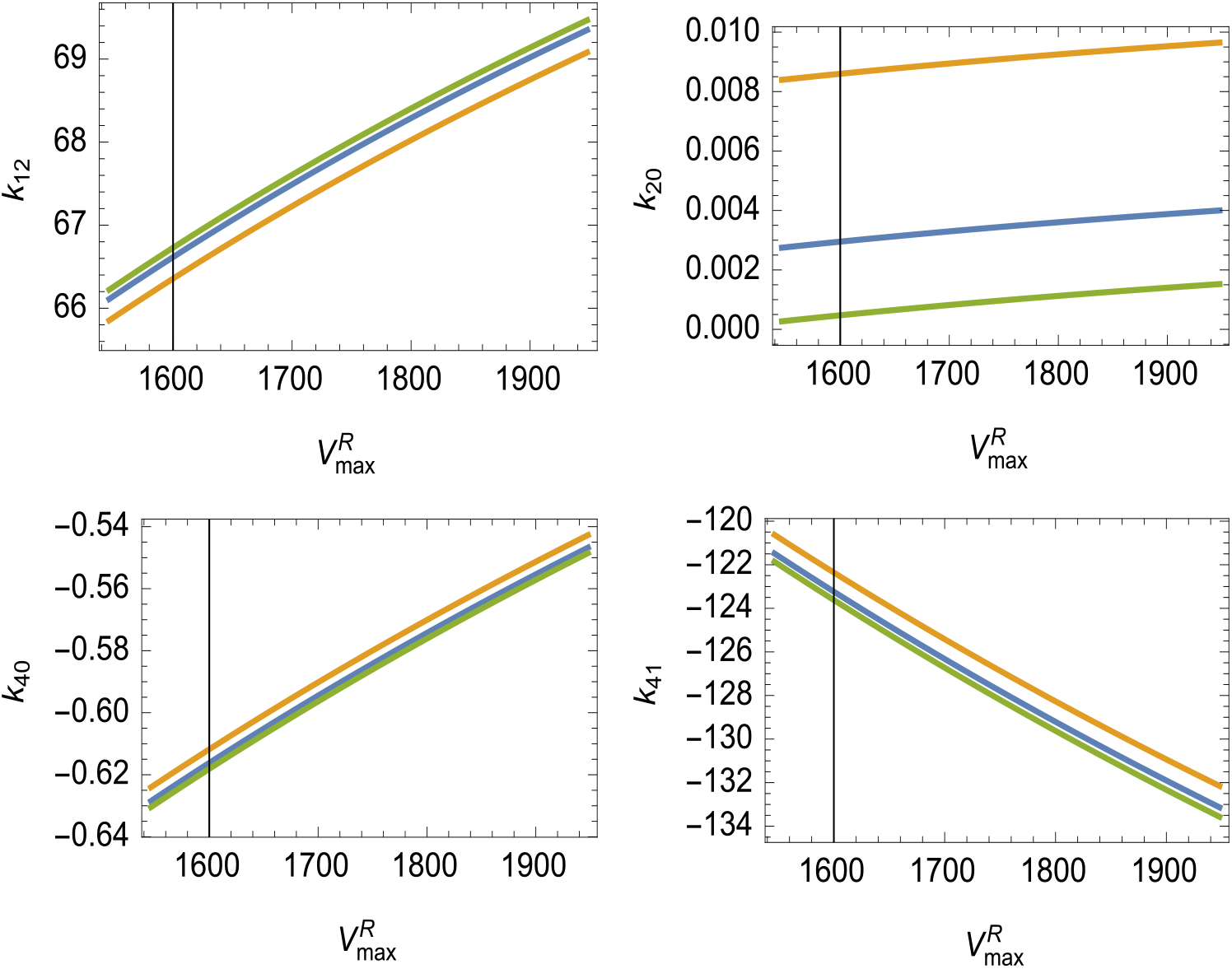
Sensitivity to the parameter 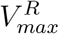 when 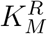 is chosen to make the Michaelis-Menten curve pass through the high (Green), low (Orange) or mid (Blue) point of the experimental error range at 1*μM* of ATP. All other parameter values were taken from Table 1.

**Figure 12:**
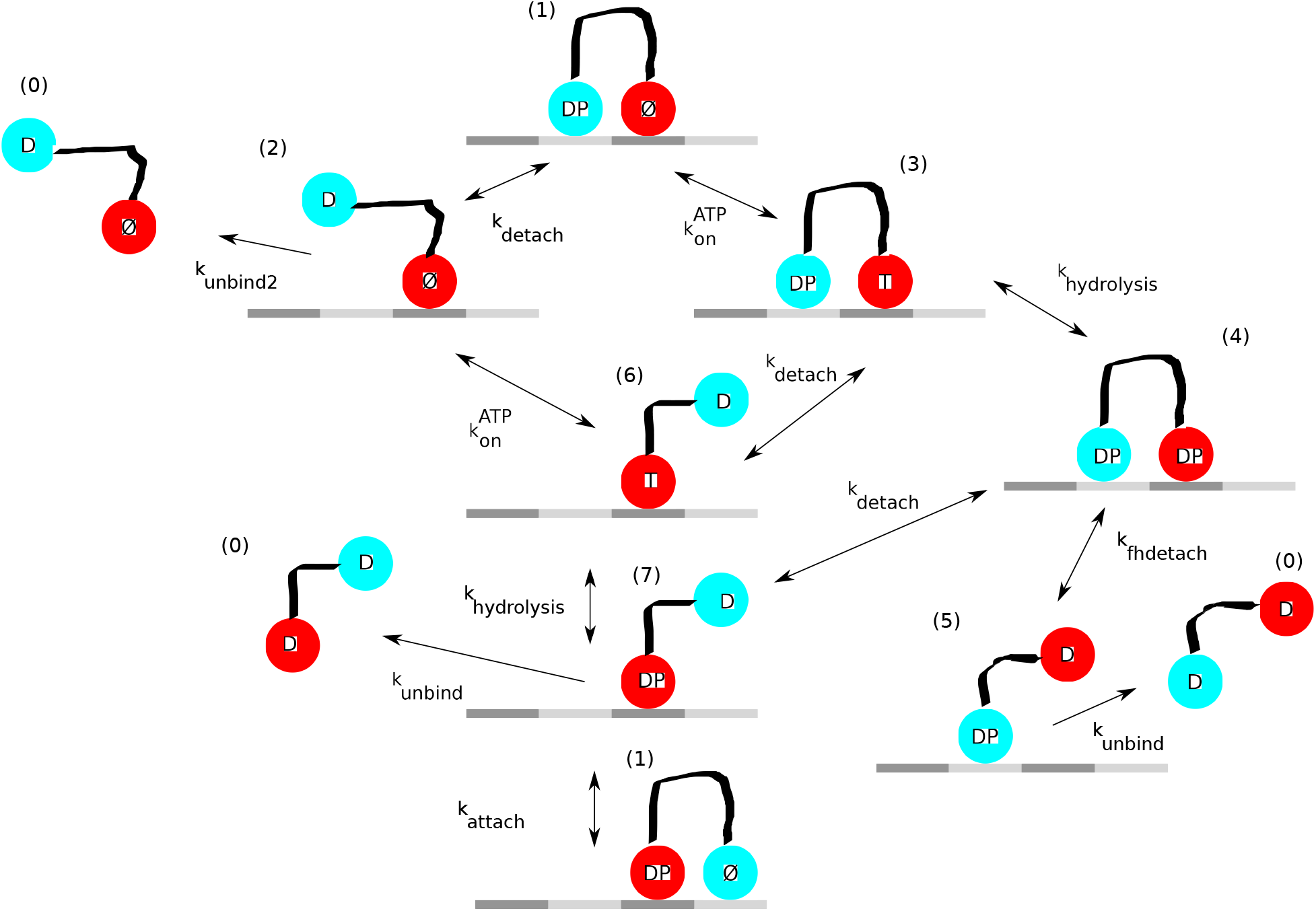
After Figure 4 from [Muthukrishnan et al., 2009]. D: ADP; T: ATP; P: Phosphate; *∅*: No nucleotide.

## 5 Kinesin-2

The model of for kinesin-2 is shown in Figure 12. For kinesin-2 the matrix for (5) is

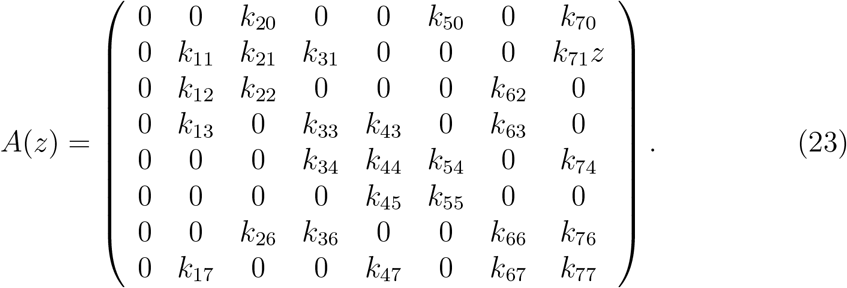

Run length (6), and run time (7) both have the Michaelis-Menten (2:2) form

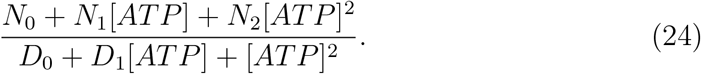

As it turns out that the denominators are the same for both exprressions, velocity also has this form. For run length, and therefore velocity, *N*_0_ = 0 for the eight parameter model in which 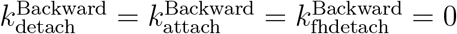.

Mathematica’s *FindFit* was successful in finding the Michaelis-Menten coefficients in (24) for run length, but failed to converge for velocity. Figure 13 shows the results of fitting curves of this form to the data for run length. If the data point at a concentration of 300 *μ*M of ATP was ignored, a good fit to the other five data points was obtained. If all six data points are used, then the curve fell outside the error bars at the upper two ATP concentrations.

**Figure 13:**
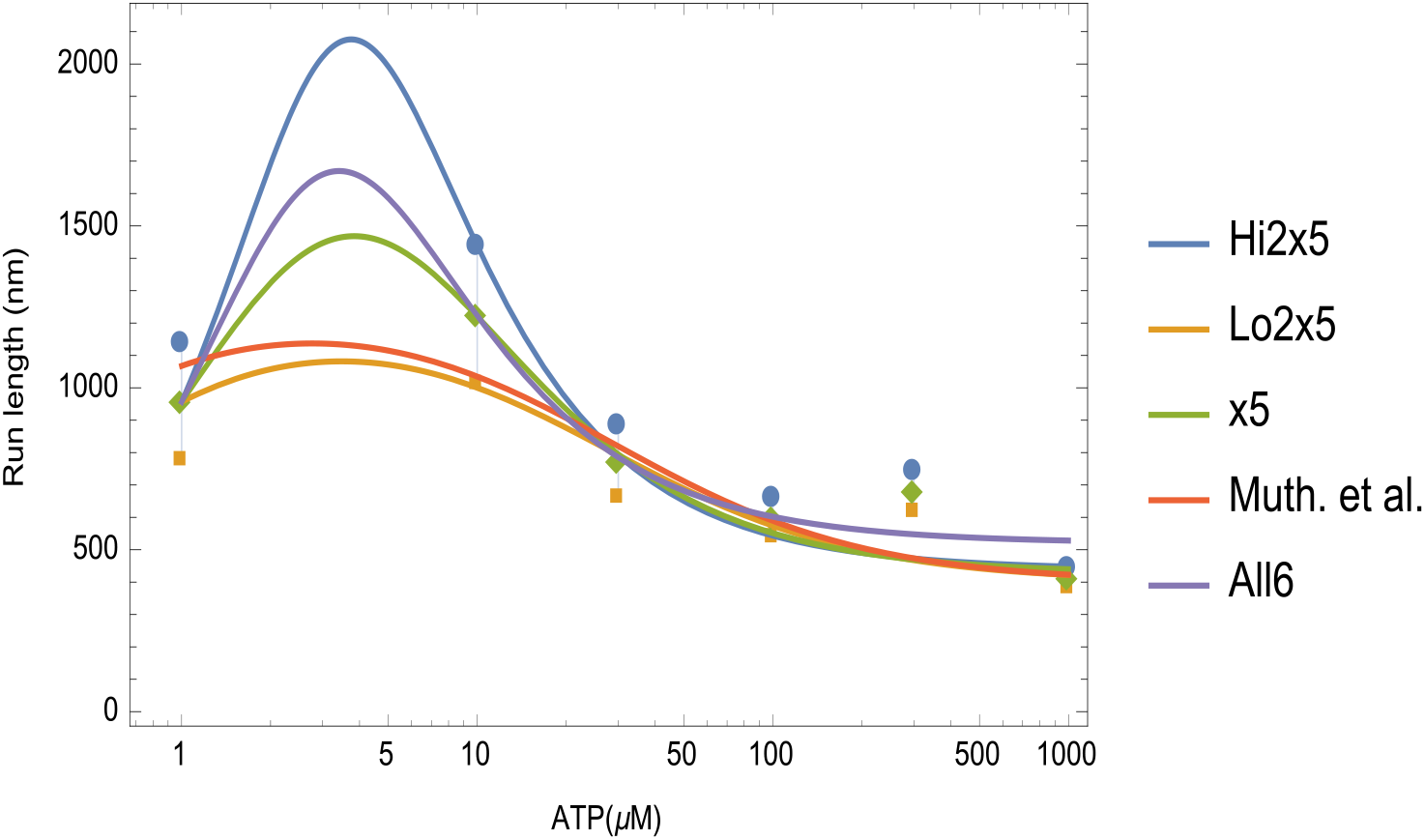
Four fittings of the kinesin-2 run length data to the form (24). Hi2×5, Lo2×5, and ×5 fit the data to the upper, lower and midpoint of the error bar for 10 *μ*M of ATP respectively, and omit the data point at 300 *μ*M of ATP. The curve All6 was the best fit to all six data points. For Muth. et al. the coefficients in (24) were calculated from the parameters in Table 3.

Attempts to solve analytically for four parameters using the Michaelis-Menten coefficients determined from experimental measurements of run length were not successful with the computing resources available if any of the backward parameters was assumed non-zero. If only the five forward and two unbinding parameters were used, it was possible to solve for four parameters in terms of the run length Michaelis-Menten coefficients and the remaing three parameters. For the results discussed here, the four parameters solved for were *k*_detach_, *k*_onATP_, *k*_hydrolysis_, and *k*_attach_. The dependence on the remaining parameters was simple in three cases: *k*_detach_ and *k*_onATP_ had the form *Constant∗k*_unbind2_, and *k*_attach_ had the form *Constant k*_unbind_. Here *Constant* depends only upon the run length Michaelis-Menten coefficients and the step size Δ*x*.The parameters *k*_hydrolysis_, *k*_fhdetach_, and *k*_unbind2_ had a nonlinear relationship.

Even if only the seven forward and unbinding parameters were retained, the problem of solving for the remaining three parameters in terms of the coefficients in the Michaelis-Menten formula for velocity was intractable. Thus numerical solution was used. The Michaelis-Menten coefficients *N*_1_*, N*_2_*, D*_0_*, D*_1_ in the form (24) for velocity were found in terms of the seven forward and unbinding parameters, and the four parameters listed above were replaced by their representations in terms of the run length Michaelis-Menten coefficients and remaining parameters. The objective function was the *l*_2_ norm of the vector of relative errors between the Michaelis-Menten velocity formula, evaluated at the candidate parameters, and the experimental data.

It turned out that this problem could not be solved numeically: the value of *k*_unbind_ retuned by *fmincon* was always its upper constraint. However if *k*_unbind_ was left as a free parameter, the other two parameters, *k*_fhdetach_ and *k*_unbind2_ could be determinined numerically. Figure 14 shows the five cases listed in Figure 13 generate a wide range of estimates for the parameters.

**Figure 14:**
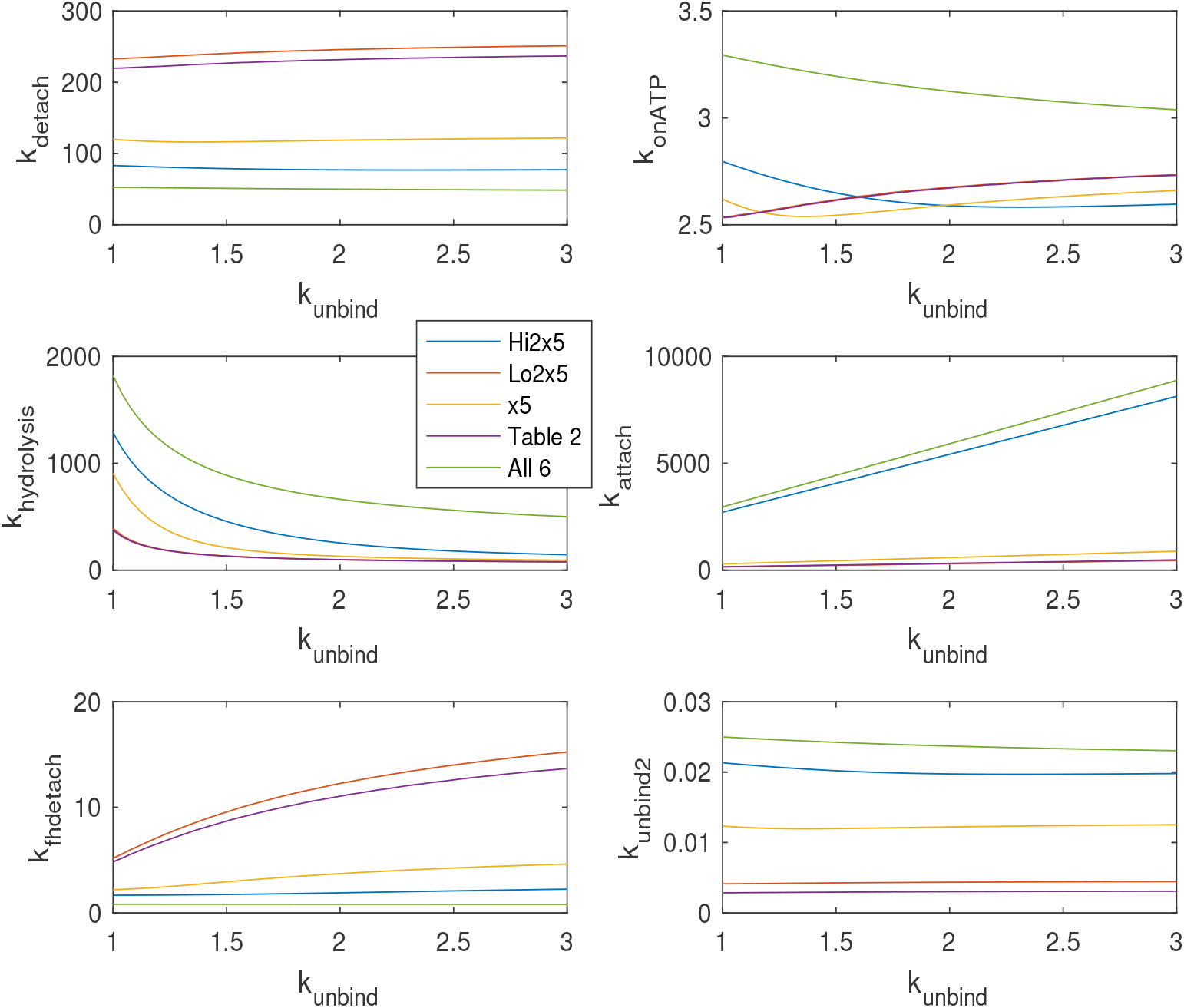
Six forward parameters of kinesin-2 as functions of *k*_unbind_.

Table 4 compares the results of the process described above with the direct global fitting of the run length and velocity data as described previously in connection with kinesin-1 (Table 2). In the global fitting cases either *k*_unbind_ or *k*_attach_ had to be determined exogenously. In the case of global fitting attempts to find one of the backward parameters met with mixed success. The backward rate *k*_hydrolysis_ was found to be about 0.001, but attempting to find the backward rate for *k*_onATP_ was destabilizing.

**Table 4:** Results of constrained minimization for kinesin-2 for the cases shown in Figure 13. The value of *k*_unbind_ was set at the value shown. For the cases labeled “G”, direct global fitting to the run length and velocity data was used. In the other cases labeled “MM” the parameter values were determined by analytically solving for four rates in terms of the Michaelis-Menten run length coefficients and then minimizing over the velocity data, as described in the text. For the “G” cases the values of 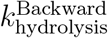 which were on the order of 0.001, are not displayed. In all cases *k*_unbind_ = 2.3 was assumed.

| Case | $k_{\text{detach}}$ | $k_{\text{onATP}}$ | $k_{\text{hydrolysis}}$ | $k_{\text{attach}}$ | $k_{\text{fhdetach}}$ | $k_{\text{unbind2}}$ | $k_{\text{unbind}}$ | Obj ftn |
| --- | --- | --- | --- | --- | --- | --- | --- | --- |
| Muth. et al. | 250 | 3 | 120 | 370 | 11 | 0.003 | 2.3 |  |
| All6G | 130 | 2.7 | 100 | 530 | 3.9 | 0.01 | 2.3 | 0.363545 |
| All6MM | 49 | 3.1 | 590 | 6800 | 0.81 | 0.023 | 2.3 | 0.481877 |
| Lo2x5G | 220 | 2.7 | 91 | 350 | 10 | 0.0043 | 2.3 | 0.197735 |
| Lo2x5MM | 210 | 2.7 | 92 | 370 | 9.6 | 0.0051 | 2.3 | 0.192171 |
| Mid2x5G | 140 | 2.6 | 100 | 570 | 5.1 | 0.011 | 2.3 | 0.228118 |
| Mid2x5MM | 91 | 2.6 | 140 | 890 | 2.4 | 0.014 | 2.3 | 0.222335 |
| Hi2x5G | 110 | 2.6 | 120 | 1200 | 3.6 | 0.016 | 2.3 | 0.258815 |
| Hi2x5MM | 71 | 2.6 | 250 | 0 | 1.7 | 0.021 | 2.3 | 0.278204 |

### 5.1 Simplified model for kinesin-2

Since the full model for kinesin-2 was analytically intractable, and there were unanswered questions about the solvability of the model, a simpler model was considered. In Figure 12, for the simplified model, the motor can unbind from state (3) at rate *k*_30_, transition to state (6) at rate *k*_36_ or to state (7) at rate *k*_37_.

Neglecting any backward transitions, the matrix (5) is

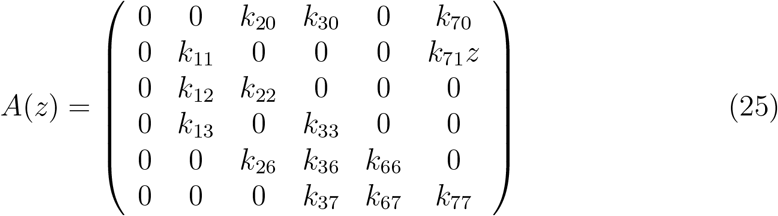

where *k*_12_ = *k*_36_ = *k*_detach_, 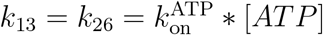, *k*_67_ = *k*_hydrolysis_, *k*_71_ = *k*_attach_, *k*_20_ = *k*_unbind2_, *k*_70_ = *k*_unbind_, the unbinding parameter *k*_30_ subsumes *k*_fhdetach_, and *k*_37_ is a new rate parameter. The simplifeid model continues to have Michaelis-Menten (2:2) form, and thus for experimentally determined values of the Michaelis-Menton coefficients there are now eight equations for eight unknown rates.

With backward transitions disallowed, run length and velocity have the form (24) with *N*_0_ = 0 as they did for the full model. The parameter *k*_hydrolysis_ was not present in the run length equation. The rates *k*_detach_, 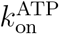, *k*_37_, and *k*_attach_ were solved for in terms of 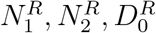, and 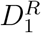 in the run length equation. results were substituted into the 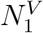 velocity equation, and *k*_hydrolysis_ was solved for. Upon substituting all five parameter formulas into the remaiining 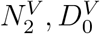 and 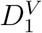 velocity equations, the result is a consistency condition

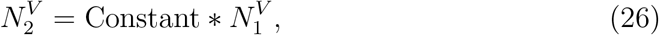

an equation

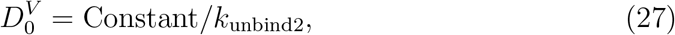

and an equation for 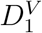 involving the remaining unknowns *k*_30_ and *k*_unbind_, after *k*_unbind2_ has been eliminated using (27). Factors designated as “Constant” depend only upon the Michaelis-Menten run length parameteers and the step length Δ*x*. Thus this simplified model of kinesin-2 determines only six paramters with the seventh being determined exogenously, as does the full model. The parameter values for the cases “Lo2×5” and “×5” are given in Table 5. Values for *k*_hydrolysis_ were lower than the corresponding cases for the full model. Values for *k*_onATP_ and *k*_unbind2_ were somewhat higher.

**Table 5:**
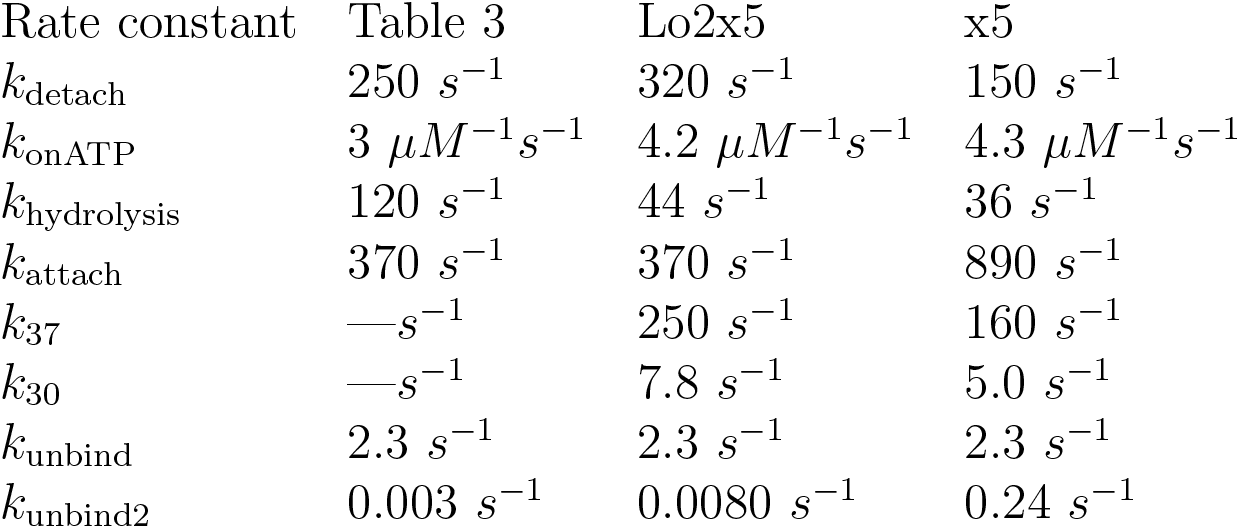
Parameters for the simplified kinesin-2 model described in the text. The rate *k*_unbind_ is assumed to be the same for all three cases.

| Rate constant | Table 3 | Lo2x5 | x5 |
| --- | --- | --- | --- |
| $k_{\text{detach}}$ | $250 \text{ s}^{-1}$ | $320 \text{ s}^{-1}$ | $150 \text{ s}^{-1}$ |
| $k_{\text{onATP}}$ | $3 \mu\text{M}^{-1}\text{s}^{-1}$ | $4.2 \mu\text{M}^{-1}\text{s}^{-1}$ | $4.3 \mu\text{M}^{-1}\text{s}^{-1}$ |
| $k_{\text{hydrolysis}}$ | $120 \text{ s}^{-1}$ | $44 \text{ s}^{-1}$ | $36 \text{ s}^{-1}$ |
| $k_{\text{attach}}$ | $370 \text{ s}^{-1}$ | $370 \text{ s}^{-1}$ | $890 \text{ s}^{-1}$ |
| $k_{37}$ | $—\text{s}^{-1}$ | $250 \text{ s}^{-1}$ | $160 \text{ s}^{-1}$ |
| $k_{30}$ | $—\text{s}^{-1}$ | $7.8 \text{ s}^{-1}$ | $5.0 \text{ s}^{-1}$ |
| $k_{\text{unbind}}$ | $2.3 \text{ s}^{-1}$ | $2.3 \text{ s}^{-1}$ | $2.3 \text{ s}^{-1}$ |
| $k_{\text{unbind2}}$ | $0.003 \text{ s}^{-1}$ | $0.0080 \text{ s}^{-1}$ | $0.24 \text{ s}^{-1}$ |

The simplified model retained some relations exhibited by the model for kinesin-1 and the full model for kinesin-2. The rate *k*_attach_ was linearly proportional to *k*_unbind_ and independent of all other rates. Similarly, the rates *k*_detach_ and *k*_onATP_ were linearly proportional to *k*_unbind2_ and independent of all other rates.

When the consistency condition (26) is substituted into the formula for velocity (24), the result is

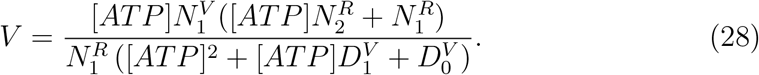

As mentioned previously for the case of (24), Mathematica’s *FindFit* was unable to find Michaelis-Menten coefficients for the velocity data. *FindFit* was, however, able to find the remaining three Michaelis-Menten velocity coefficients in (28) for given run length coefficients. See Figure 15.

**Figure 15:**
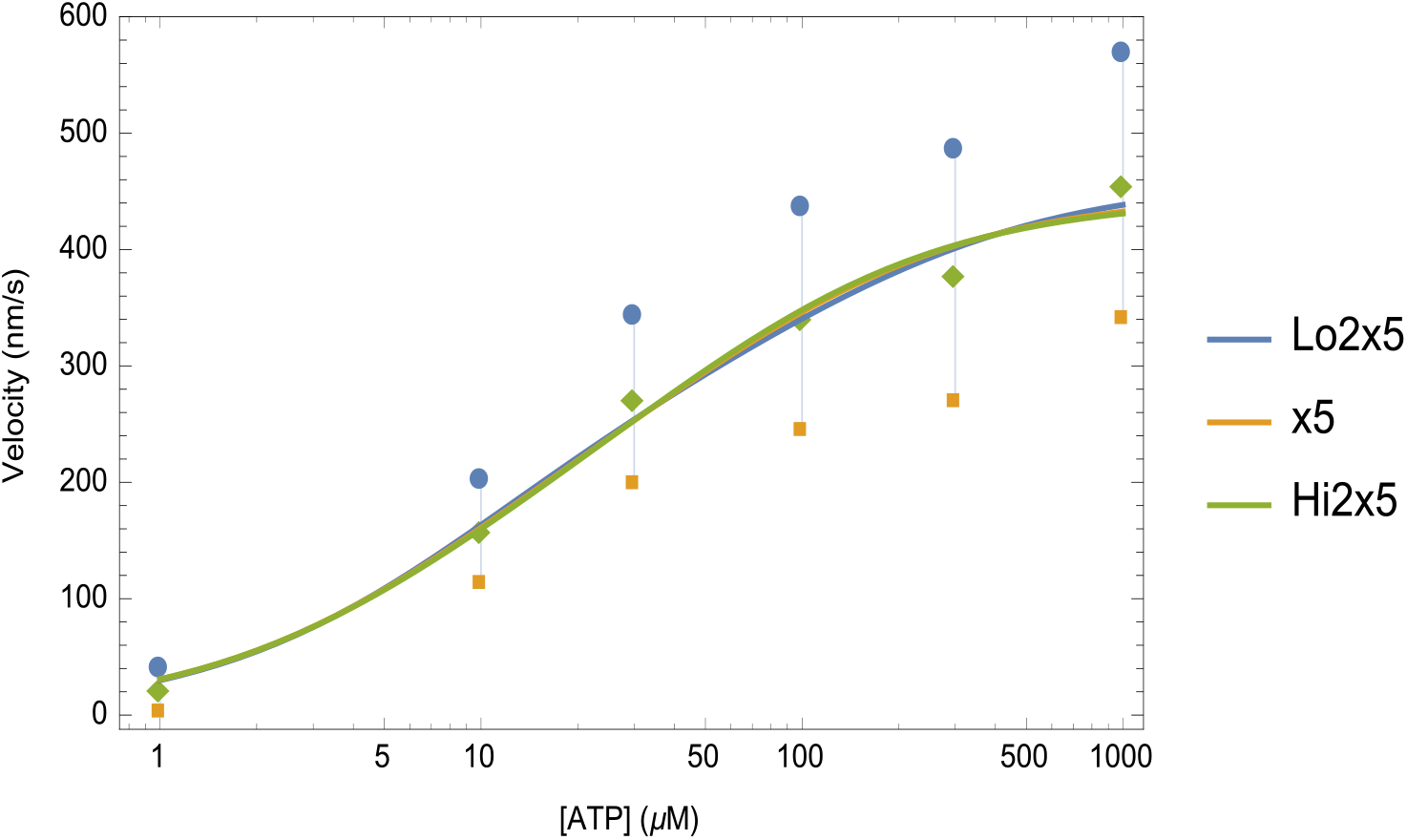
Velocity calculated from equation (28) for three of the cases in Figure 13.

## 6 Myosin VI

The article [Elting et al., 2011] gives experimental data on run length and velocity for two varieties of myosin VI, a wild type and an engineered form, which will be designated types I and II.

The chemo-mechanical cycle shown in Figure 16, leads to

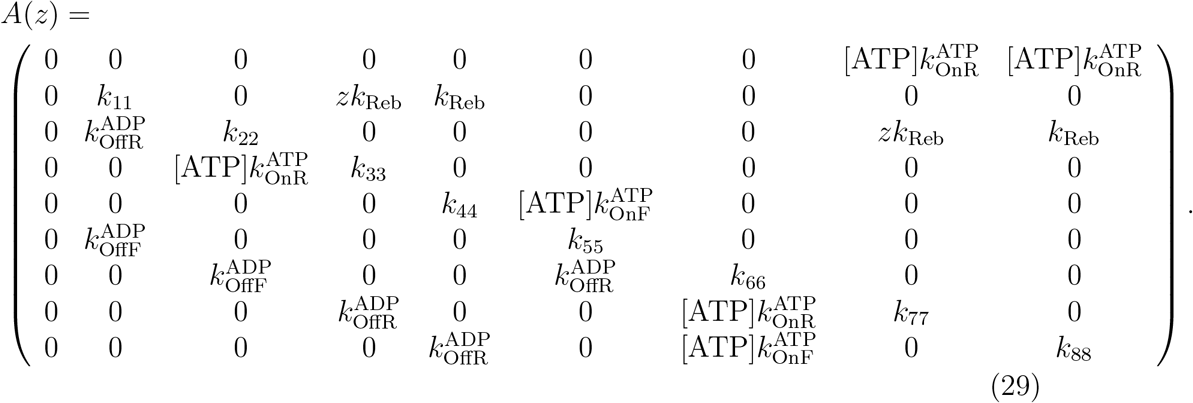

**Figure 16:**
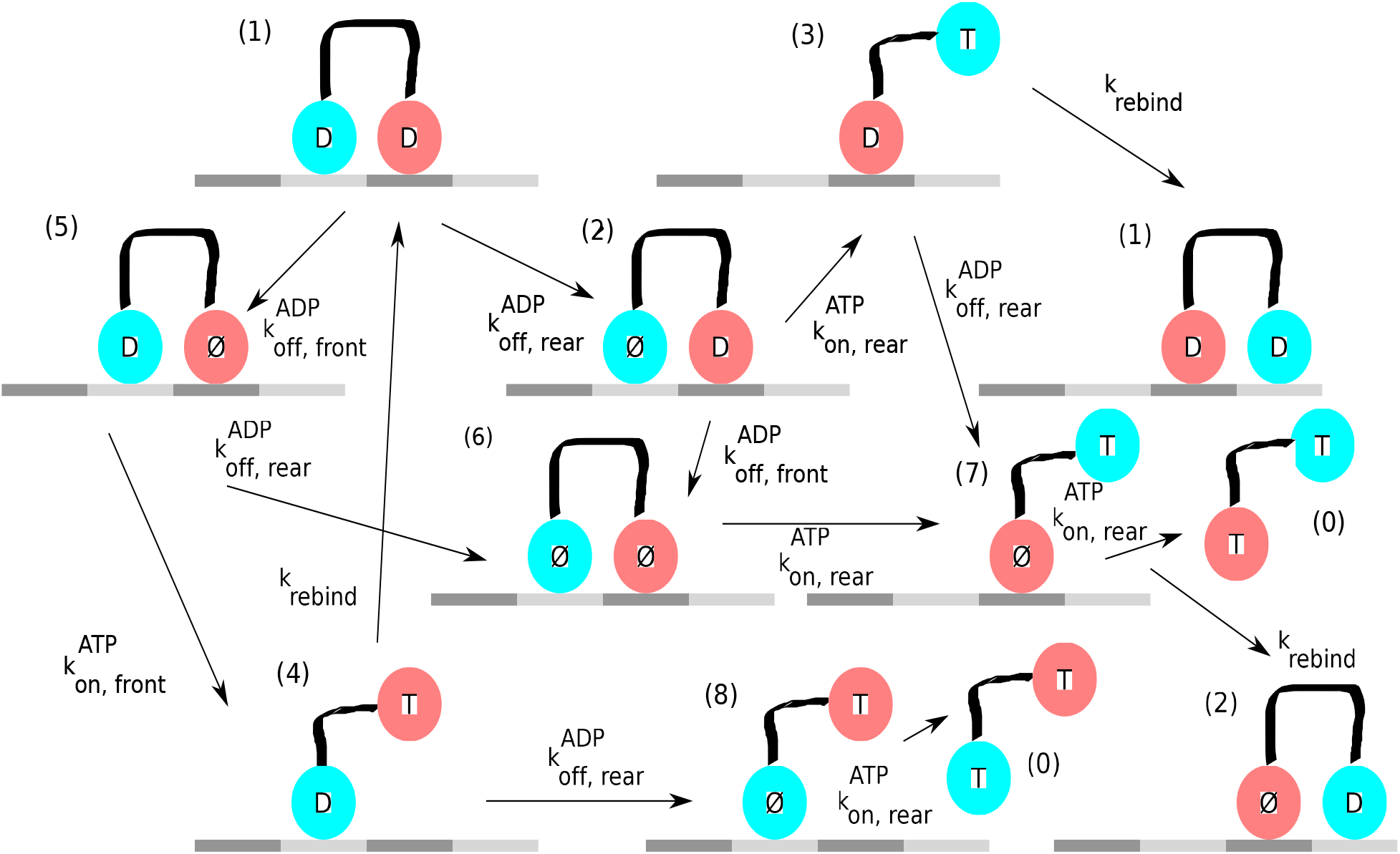
After Figure 2 from [Elting et al., 2011]

The resulting run length and run time formulas have Michaelis-Menten (4:4) forms, but velocity has a (5:5) form. Figure 17 shows that the parameters in Table did not give good agreement with the experimental measurements in [Elting et al., 2011], when the parameters of Table 6 were used.. As noted in the article defects in the actin filament may have led to premature detachments.

**Figure 17:**
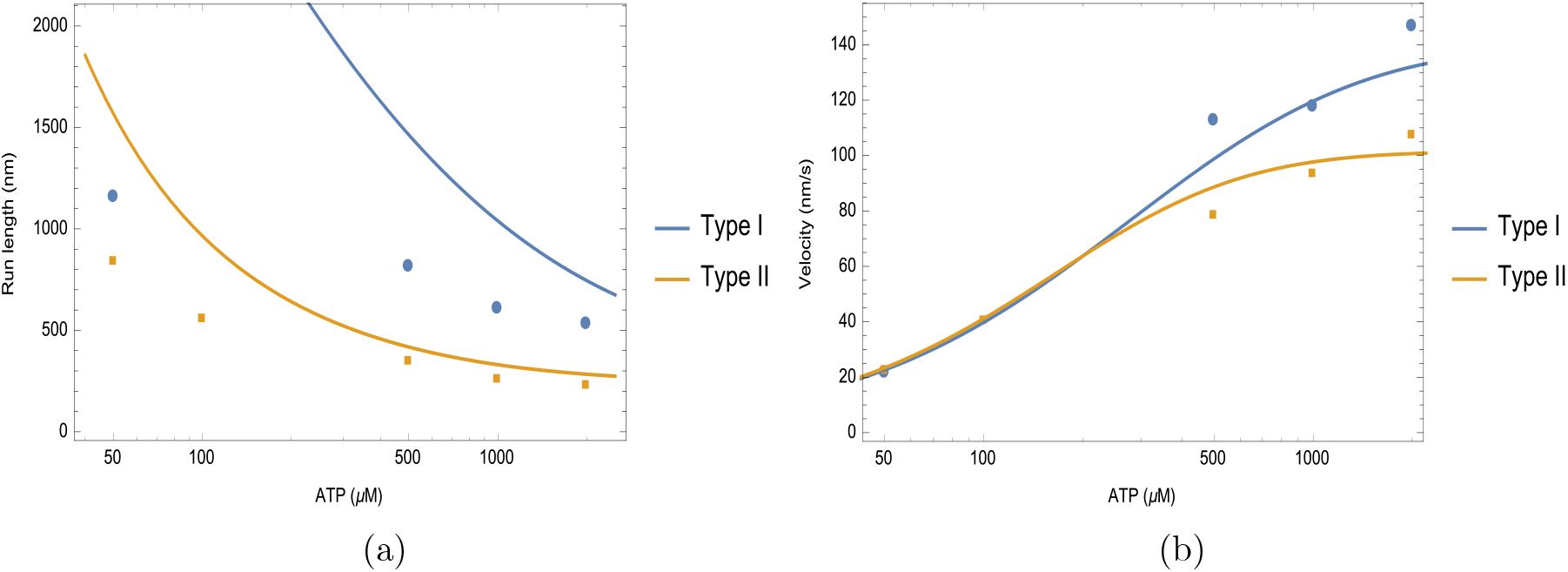
Predictions of the Markov model for two types of myosin VI. The parameters used are given in Table 6. The experimental data was read from Figure 3 of [Elting et al., 2011].

**Table 6:**
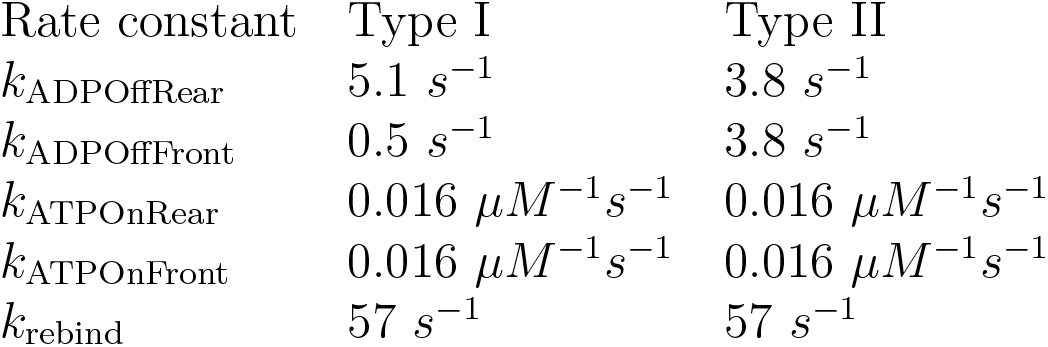
Parameters for the myosin VI model of Figure 16 from [Elting et al., 2011]. Type I is a wild-type myosin VI, Type II an engineered vari-ety.

| Rate constant | Type I | Type II |
| --- | --- | --- |
| $k_{\text{ADPOffRear}}$ | $5.1 \text{ s}^{-1}$ | $3.8 \text{ s}^{-1}$ |
| $k_{\text{ADPOffFront}}$ | $0.5 \text{ s}^{-1}$ | $3.8 \text{ s}^{-1}$ |
| $k_{\text{ATPOnRear}}$ | $0.016 \mu\text{M}^{-1}\text{s}^{-1}$ | $0.016 \mu\text{M}^{-1}\text{s}^{-1}$ |
| $k_{\text{ATPOnFront}}$ | $0.016 \mu\text{M}^{-1}\text{s}^{-1}$ | $0.016 \mu\text{M}^{-1}\text{s}^{-1}$ |
| $k_{\text{rebind}}$ | $57 \text{ s}^{-1}$ | $57 \text{ s}^{-1}$ |

The presence of defects may be modeled approximately by adding an unbinding rate *k*_Unbind_ at the vunerable states (3), (4), (7) and (8) in Figure 16 where only one head is attached. Figure 18 shows that better agreement with the experimental data is obtained with *k*_Unbind_ = 1.1 s^−1^. However this is an ad hoc adjustment, and the case of myosin VI was not pursued further.

**Figure 18:**
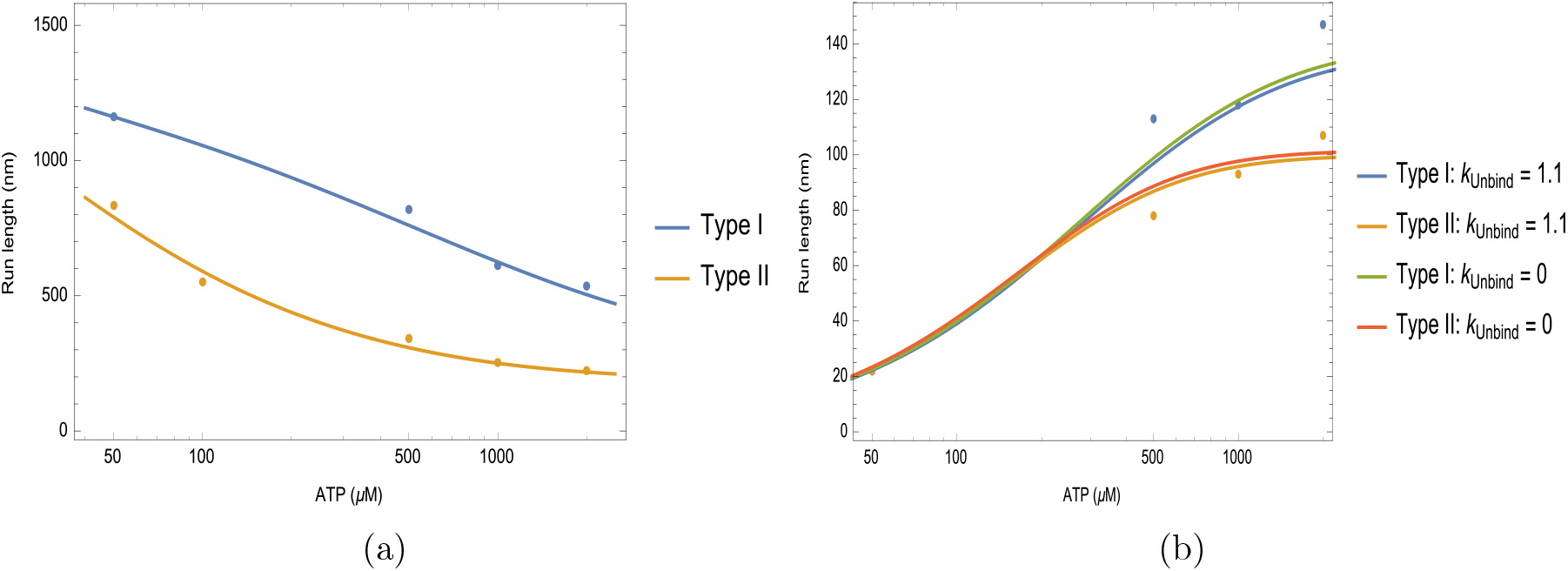
Revised predictions of the Markov model for two types of myosin VI when allowance is made for unbinding in vunerable states due to defects in the actin filaments.

## 7 Summary

The authors of [Muthukrishnan et al., 2009] and [Elting et al., 2011] took different approaches to mathematically modeling the protein motors they considered. In this article we have shown that a Markov modeling approach fits all the chemo-mechanical cycles considered. In all cases both run length and run time had Michaelis-Menten forms of orders that increased with the complexity of chemo-mechanical cycle. Due to cancellations, velocity had a lower order Michaelis-Menten form than the quotient of run length and run time would suggest.

Four models were considered. A simple three stage model illustates that al-lowing for both forward and backward stepping roughly doubles the number of Markov states required.

For the four stage model of kinesin-1 presented in [Muthukrishnan et al., 2009] shown in Figure 4, run length and velocity had conventional Michaelis-Menten (1:1) form if backward transition from state (4) to (1) was disallowed. Thus at most four reaction coefficients can be found from a determination of the four Michaelis-Menten coefficients from experimental data. Direct global fitting of the full run length and velocity formulas to experimental measurements seemed to confirm that no more than four reaction rates were attainable in practice, even though there were six Michaelis-Menten coefficients in the full formulas (13) and (14), if backward transition was allowed.

Table 2 shows that not all subsets of four parameters could be obtained by direct global fitting to run length and velocity data. Analysis of the Michaelis-Menten coefficients showed that this could be due to inconsistencies in the equations, extreme sensitivity of the dependence of one parameter on another due to the presence of vertical asymptotes, or negative reaction coefficient values. When solution for a quadruplet of reaction coefficients was both analytically and numerically feasible, the parameter values were in agreement with the estimates of [Muthukrishnan et al., 2009]. With the exception of the detachment parameter *k*_unbind2_, reaction rates exhibited little sensitivity to the magnitude of measurement errors indicated by the SE bounds (Figures 8 and 9).

For the more complicated chemo-mechanical models of kinesin-2 and myosin VI, while it was possible to show that they had higher order Michaelis-Menten form and express the coefficients in terms of the reaction rates in the model, it was not possible to solve fully for the reaction rates in terms of the Michaelis-Menten coefficients, at least with the computing resources available. Thus at least some parameters would have to be estimated numerically.

For kinesin-2 run length and velocity had Michelis-Menten (2:2) form, meaning that it might be possible to solve for up to eight reaction rates. Table 4 shows that in general the two approaches of global fitting of the run length and velocity data and the hybrid analytic-numeric described in the text may lead to very different estimates for the parameters. However for the case of “Lo2×5” of Figure 13, which produced the lowest objective function values, the results of the two approaches gave similar estimates and these were for the most part consistent with the estii-mates of [Muthukrishnan et al., 2009]. Unlike the case of kinesin-1 (Figures 8 and 9), Table 4 and Figure 14 show that a wide range of parameter values are compatible with the experimental measurements for kinesin-2 and their SE estimates. Knowlwdge of the Michaelis-Menten form of run length and velocity may help in evaluating the experimental data. For instance for kinesin-1 Figure 6 suggests the use of all data points is compatible with the derived MM form, but for kinesin-2 Figure 13 suggests that the fifth data point is not compatible with the other five and the derived Michaelis-Menten (2:2) form.

The third molecular motor considered, myosin VI, had the most complicated chemo-mechenical cycle resulting in (4:4) and (5:5) Michaelis-Menten forms for run length and velocity, respectively. The predictions did not match the experimental measurements in [Elting et al., 2011], possibly because the presence of defects in the actin filaments leading to premature detachment.

## A Appendix Expected run length and run time

Following [Elston, 2000] define the moment generating function

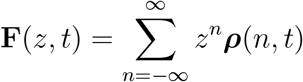

and note the following properties

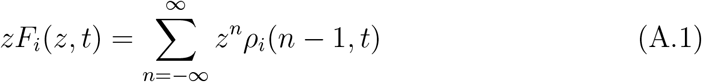

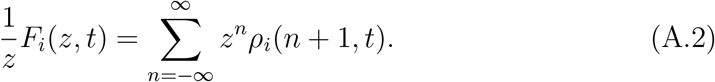

The first two moments of the random variable *N* (*t*), the number of steps taken, are

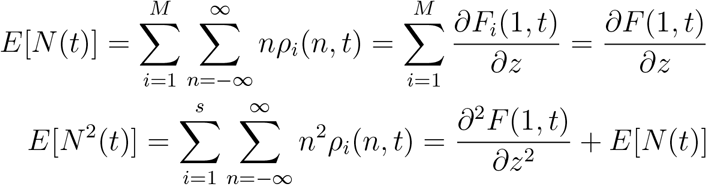

where *M* is the number of non-absorbing states, and 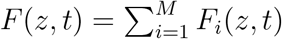. From (2), (A.1), and (A.2), the generating function satisfies the differential equation

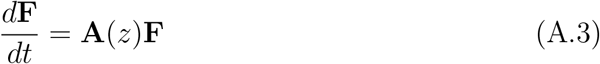

where **A** is given by (5).

Let the first column of **A**(*z*) represent detachment. For *z* near 1, *λ*_0_(*z*) ≡ 0 is the largest eigenvalue with right eigenvector **e**^(0)^ = [1, 0, …, 0]^*T*^. Assume Re*λ*_*M*_ < Re*λ*_*M* − 1_ < … < Re*λ*_1_ < 0. The solution to (A.3) corresponding to the initial condition **F**(*z*, 0) = **p**^(0)^ = [0, *p*_1_, …, *p*_*M*_]^*T*^ is

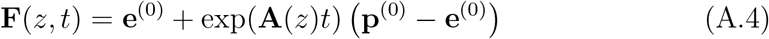

Let **S** be the matrix for which

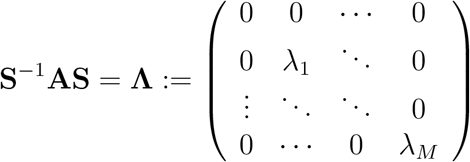

The columns and rows of **S** and **S**^−1^ are the right and left eigenvectors of **A**(*z*), respectively. Thus the first column of **S** is **e**^(0)^, whence the first column of **S**^−1^ is also **e**^(0)^. From (A.4)

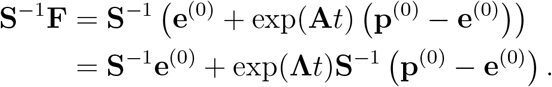

Let **r**_*j*_, *j* = 0, …, *n* − 1 denote the columns of **S** and **l**_*j*_, *j* = 0 …, *n*−1 the rows of **S**^−1^. Then

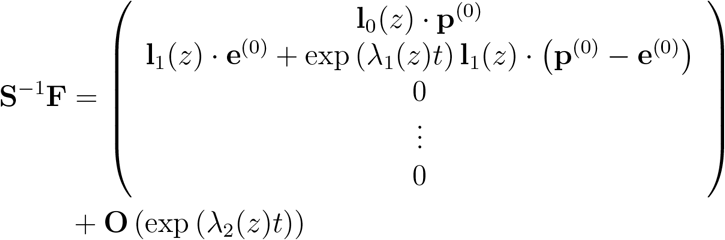

Since **r**_0_(*z*) *≡* [1, 0, …, 0]^*T*^, and **l**_1_(*z*) *·* **e**^(0)^ = 0,

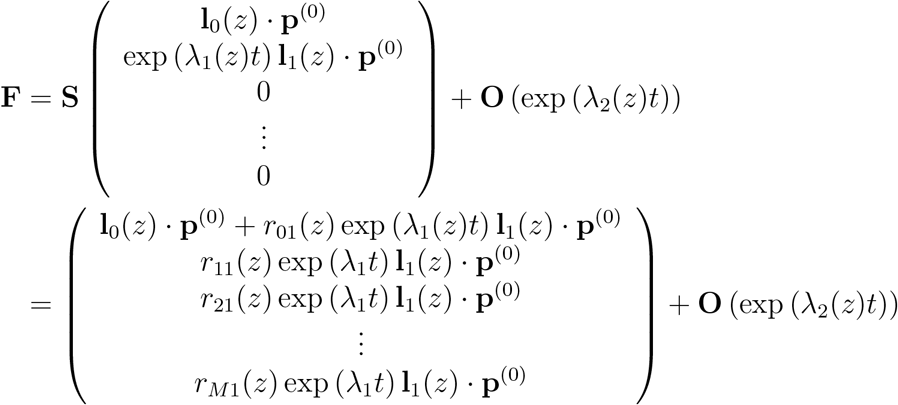

Thus

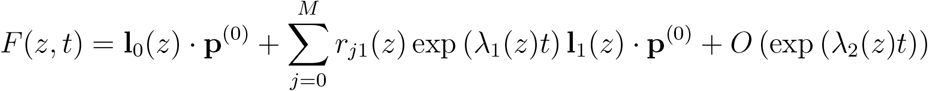

and

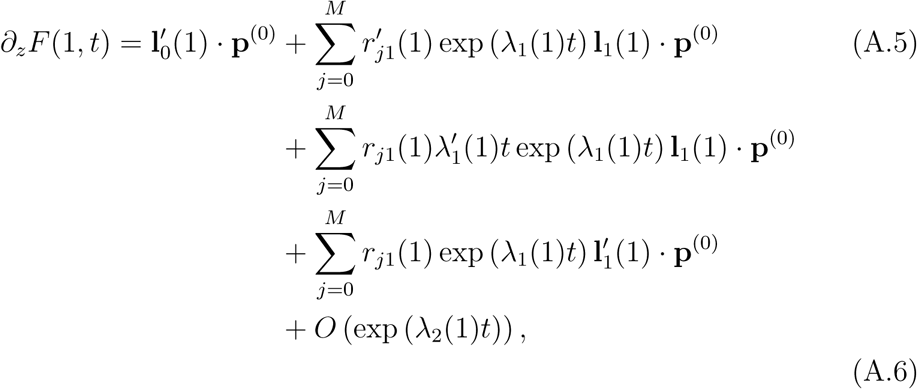

whence 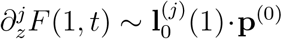 and for a molecular motor that takes steps of length Δ*x*, the asymptotic expected run length and its variance are

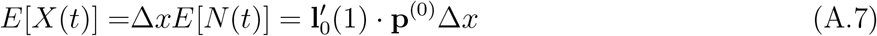

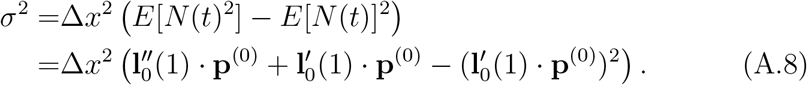

The matrix **A**(1) given by (5) represents a continuous-time Markov process. Hence the formula for the random variable *T* representing time to detachment is a phase-type distribution with moments

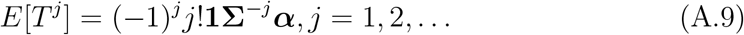

where **1** is a row vector of 1’s,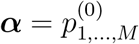 and the subgenerator matrix for this problem is **Σ** = *A*(1)_1,…,*M*,1,…,*M*_.

## References

Asbury, C. L., Fehr, A. N., and Block, S. M. (2003). Kinesin moves by an asymmetric hand-over-hand mechanism. Science, 302(5653):2130–2134.

Elston, T. C. (2000). A macroscopic description of biomolecular transport. Journal of Mathematical Biology, 41(3):189–206.

Elting, M. W., Bryant, Z., Liao, J.-C., and Spudich, J. A. (2011). Detailed tuning of structure and intramolecular communication are dispensable for processive motion of myosin VI. Biophysical Journal, 100(2):430–439.

Gillespie, D. T. (1977). Exact stochastic simulation of coupled chemical reactions. The journal of physical chemistry, 81(25):2340–2361.

Hancock, W. O. and Howard, J. (1999). Kinesin’s processivity results from mechanical and chemical coordination between the atp hydrolysis cycles of the two motor domains. Proceedings of the National Academy of Sciences, 96(23):13147–13152.

Muthukrishnan, G., Zhang, Y., Shastry, S., and Hancock, W. O. (2009). The processivity of kinesin-2 motors suggests diminished front-head gating. Current Biology, 19(5):442–447.

Ross, J. L., Ali, M. Y., and Warshaw, D. M. (2008). Cargo transport: molecular motors navigate a complex cytoskeleton. Current Opinion in Cell Biology, 20(1):41–47. ¡ce:title¿Cell structure and dynamics¡/ce:title¿.

Shastry, S. and Hancock, W. O. (2010). Neck linker length determines the degree of processivity in kinesin-1 and kinesin-2 motors. Current Biology, 20(10):939–943.

Yajima, J., Alonso, M. C., Cross, R. A., and Toyoshima, Y. Y. (2002). Direct long-term observation of kinesin processivity at low load. Current Biology, 12(4):301–306.

